# Evidence for evolution of a new sex chromosome within the Marchantiales haploid-dominant plant lineage

**DOI:** 10.1101/2025.01.16.633329

**Authors:** Yuan Fu, Xiaoxia Zhang, Tian Zhang, Wenjing Sun, Wenjun Yang, Yajing Shi, Jian Zhang, Qiang He, Deborah Charlesworth, Yuannian Jiao, Zhiduan Chen, Bo Xu

**Affiliations:** State Key Laboratory of Plant Diversity and Prominent Crop/State Key Laboratory of Systematic and Evolutionary Botany, Institute of Botany, Chinese Academy of Sciences, Beijing 100093, China; China National Botanical Garden, Beijing 100093, China; University of Chinese Academy of Sciences, Beijing 100049, China; Institute of Evolutionary Biology, University of Edinburgh, Edinburgh EH9 3FL, UK

## Abstract

Sex chromosomes have evolved independently in numerous lineages across the tree of life, in both diploid-dominant species, including many animals and plants, and the less studied haploid-dominant plants and algae. Strict genetic sex determination ensures that individuals reproduce by outcrossing. However, species with separate sexes (termed dioecy in diploid plants, and dioicy in haploid plants) may sometimes evolve different sex systems, and become monoicous, with the ability to self-fertilize. Here, we study dioicy-monoicy transitions in the ancient liverwort haploid plant lineage, using three telomere-to-telomere gapless chromosome-scale reference genome assemblies from the Ricciaceae group of Marchantiales. Ancestral liverworts are believed to have been dioicous, with U and V chromosomes (chromosome 9) determining femaleness and maleness, respectively. We confirm the finding that monoicy in *Ricciocarpos natans* evolved from a dioicous ancestor, and most ancestrally U chromosomal genes have been retained on autosomes in this species. We also describe evidence suggesting the possible re-evolution of dioicy in the genus *Riccia*, with probable *de novo* establishment of a sex chromosome from an autosome (chromosome 5), and further translocations of genes from the new sex chromosome to autosomes. Our results also indicated that m-chromosomes are consistent genomic features, and may have evolved independently from sex chromosomes in *Ricciocarpos* and *Riccia* lineages.

## Introduction

Genetic sex determining systems involving XY or ZW sex chromosome systems, and their evolution, have been intensively studied in organisms with dominant diploid life stages, including mammals (Cortez et al., 2014), insects (Toups and Vicoso, 2023), birds (e.g. Sigeman et al., 2021), fish (El Taher et al., 2021; Xue et al., 2021), and seed plants (Carey et al., 2021a; Charlesworth and Harkess, 2024). Much less studied are the diverse lineages of haploid organisms, including bryophytes and green and brown algae, though it has long been known that their haploid gametophyte stages may show separate sexes controlled by sex-determining regions of sex chromosomes termed U for the female- and V for the male-determining chromosomes (reviewed in Bachtrog et al., 2011; Coelho et al., 2018; Carey et al., 2021b; Iwasaki et al., 2021; Singh et al., 2023). At least some UV systems are ancient, possibly older than any known XY and ZW systems, but few genomes have currently been sequenced, so neither their homologies, nor the sometimes giant or sometimes minute sizes of their U and/or V chromosomes are currently well understood (Allen, 1917; Anderson, 1984; Renner et al., 2017).

The liverworts, one major bryophyte lineage, include more than 7,000 extant species distributed all over the world (Laenen et al., 2015; Wang et al., 2022). The Marchantiales from the second largest liverwort lineage, and are estimated to have evolved over 196 million years ago (mya) (Villarreal et al., 2016). They have adapted to habitats ranging from terrestrial to aquatic environments (Wang et al., 2022). Changes in mating systems often accompany habitat changes. For example, species that adapt to ephemeral aquatic environments, or colonizing species, often evolve the ability to reproduce in the absence of conspecifics through losing separate sexes (Carlquist, 1966), an example of “Baker’s law” (Baker, 1955). Independent transitions from dioicy to monoicy have evolved in numerous liverwort clades (Wang et al., 2022; Singh et al., 2023). The first complete liverwort genome sequenced was that of a dioicous species, *Marchantia polymorpha* (Bowman et al., 2017). Transitions from dioicy to monoicy have been suggested within the Marchantiales (see Fig. 1; Table S1), including the evolution of monoicy in *Ricciocarpos natans* from a dioicous ancestor (Singh et al. 2023). Sequencing genomes of haploid plant species from more lineages will offer opportunities to study changes in the sex chromosomes during such changes between different sexual systems.

**Figure 1.**
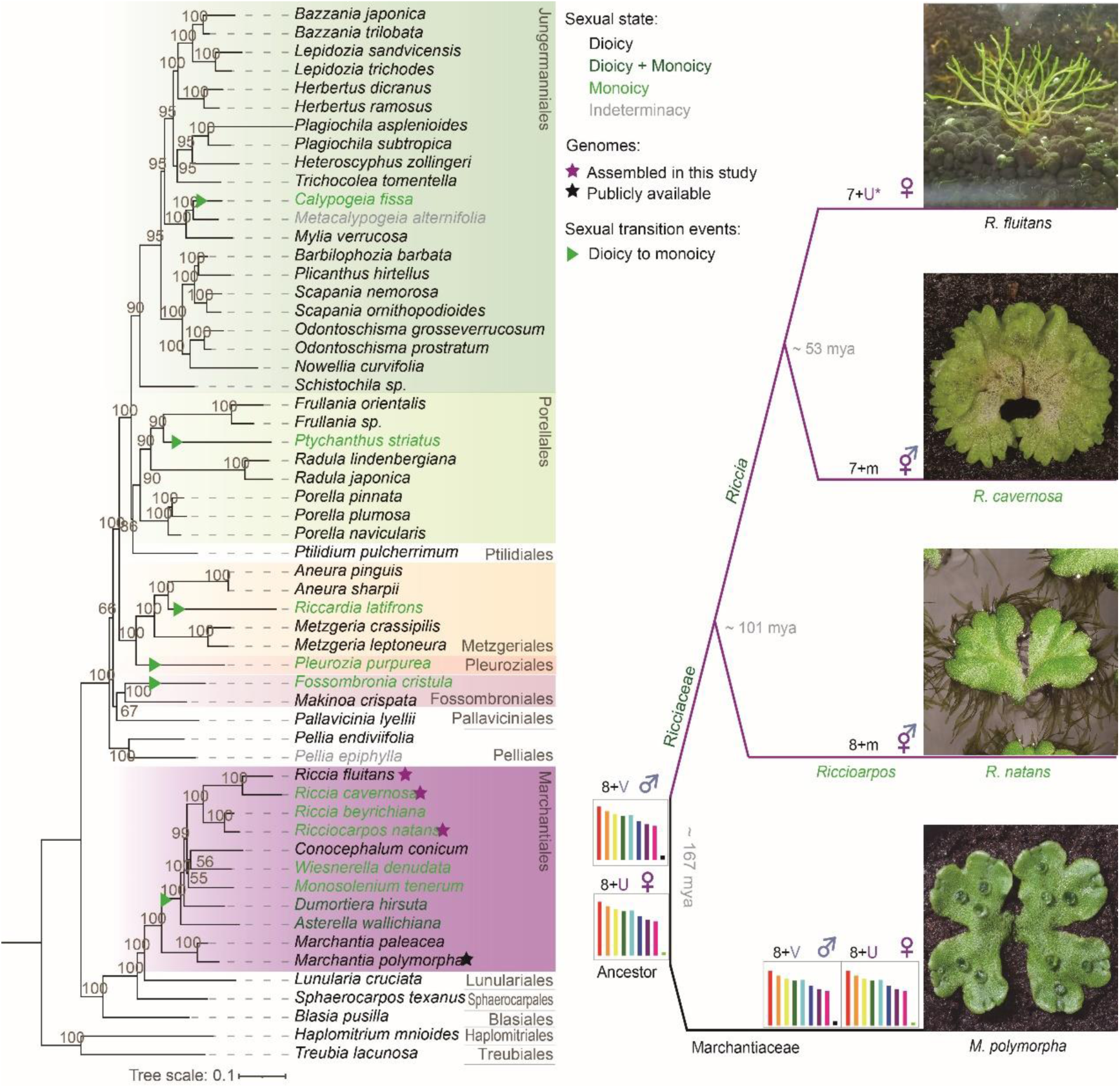
Phylogeny of the Marchantiophyta and karyotypes of the four liverwort species studied here. The phylogeny at the left is based on conserved sites in 446 low copy-number genes identified from bryophyte transcriptome sequences (as detailed in the Methods section). The figure illustrates independent sexual transitions inferred in different Marchantiophyta lineages (the different colored boxes); these interpretations are based on the species included in this analysis, and further undetected changes might be revealed if further species were studied. Six putative transitions from ancestral dioicy to monoicy (indicated by green arrowheads on the branches) are inferred when monoicous species form a minority in a predominantly dioicous group of species. Species in black, dark green, green, and gray, respectively indicate the following sexual system: dioicy, both dioicy and monoicy, monoicy, and indeterminate. Stars indicate species for which a genome dataset was generated (purple stars) or publicly available data and used (black stars) in the present study. Karyotypes of the four species investigated are displayed at the right. Divergence times for these four species were estimated using the publicly available *M. polymorpha* genome sequence and the three genome assemblies generated in this study (see Methods), and are indicated in the right-hand part of the figure (mya indicates millions of years before the present) (Villarreal et al., 2016). The possible present or former U chromosome is indicated by U*.

The ancestral sexual system of the Marchantiales is thought to be dioicy (Berrie, 1963; Bowman, 2016), but (as mentioned above) one or more transitions from dioicy to monoicy have occurred during the evolution of the Ricciaceae lineage since its divergence from the distant Marchantiaceae lineage, which includes *M. polymorpha*. One genus, *Ricciocarpos*, includes only the monoicous *R. natans*, but there are more than 150 *Riccia* species, many of which are monoicous (Allen, 1935; Smith, 1979; Na-Thalang, 1980; Gradstein, 2017; Premita Devi et al., 2019), though dioicy has also been reliably reported (McAllister, 1928).

Here, we studied two *Riccia* species, *R. fluitans* and *R. cavernosa*, and an outgroup species (*R. natans*, see above); a phylogeny constructed using 446 low copy-number genes from 57 bryophyte species confirms that *Ricciocarpos* is an outgroup to *Riccia* (Fig. 1; Table S1; Figs. S1 and S2). These three species have different sexual systems. Like *R. natans*, *Riccia cavernosa* is monoicous and produces abundant antheridia and archegonia (The World Flora online; Figs. S2). However, *R. fluitans* and its polyploid form, *R. rhenana* (Wyatt and Davison, 2013, ÖZenoĞLu et al., 2019), have been described as dioicous. *R. fluitans*, one of the two species studied here, is not monoicous. Females have repeatedly been observed (Paton, 1973; Wyatt and Davison, 2013), but only a single report found plants with sporophytes (Paton, 1973), and individuals with antheridia have not been reported in the wild, or in experiments aimed at discovering potential sex-inducing conditions (Berrie, 1964; Paton, 1973; Althoff and Zachgo, 2020); the species is therefore described as dioicous, not monoicous. However, it may reproduce by female-only vegetative reproduction, as is common in bryophytes (reviewed in Longton, 1976 and Laenen et al., 2015), and could be wholly or largely unable to produce antheridia. Loss of sexual reproduction is predicted to evolve under conditions of low density or rarity of conspecifics, especially in species with outcrossing systems such as dioicy (Charlesworth, 1980). Most such evolutionary changes are recent, because long-term persistence is unlikely in the absence of recombination (Schwander and Crespi, 2009). Consistent with *R. fluitans* having recently changed to vegetative reproduction, another member of the *R. fluitans* complex, *R. stenophylla* (Wyatt and Davison, 2013), is monoicous. The *R. fluitans* female studied allows us to study a *Riccia* U chromosome from this lineage; currently, no V chromosome is available from this lineage, but below we describe evidence suggesting that *R. cavernosa* may carry a V.

## Results

### Genome sequences

To study the genomic and chromosomal changes involved in sexual system differences between the three Ricciaceae species, sterile culture isolates from single colonies of all species were established and subjected to PacBio long-read sequencing and high-throughput chromatin conformation capture (Hi-C) analysis to generate chromosome-scale *de novo* assemblies (Table S2; Fig. S3). The sequences were assigned to the known numbers of chromosomes, 9 in the outgroup species, *R. natans* (Siler, 1934), and one less in *R. cavernosa* (Na-Thalang, 1980), and *R. fluitans* (Wyatt and Davison, 2013) (Fig. 1; Table S3); the respective assembly lengths were 204Mb, 233Mb, and 450Mb. The completeness values of these assembled genomes, tested by Benchmarking Universal Single-Copy Orthologs (BUSCO), were 98.5%, 96.1%, and 96.8%, respectively. The genome assembly continuity was also evaluated using long-terminal-repeat retrotransposons (LRT-RTs) (Ou and Jiang, 2018). LTR assembly indexes (LAI: 10.64, 11.13, and 13.83) of *R. natans*, *R. cavernosa*, and *R. fluitans* are all higher than those of the *M. polymorpha* reference genome v6.1 (BUSCO: 95.3%; LAI: 9.2; Table S3.1).

Pairwise synonymous site divergence values within the four assembled genomes showed low values, and no evidence of whole-genome duplications in any of these species (Fig. S4), unlike mosses (Gao et al., 2020). The genome size of *R. fluitans* is almost double those of the other liverwort species, including *M. polymorpha*, due to higher repeat content; the estimated genome-wide transposable element density is 58%, versus less than 38% for the other species (Table S3.1). Using RNA iso-sequencing (PacBio HiFi) and short-read sequencing (PE150), we annotated similar numbers of protein-coding genes in all three species (15,583, 20,200, and 20,308 in *R. natans*, *R. cavernosa*, and *R. fluitans*, respectively), with BUSCO scores of 99.2%, 98.8%, and 98.8%, respectively (Table 3.1). The gene numbers are similar to the number of ∼20,000 for *M. polymorpha*, although some gene families may be smaller in *R. natans* (Fig. S5; Table S4). The assemblies of most chromosomes included telomere sequences, making them the first reported complete liverwort telomere-to-telomere gap-free chromosome-scale genome assemblies with high quality and accuracy (Fig. 2A; Table S3.2).

**Figure 2.**
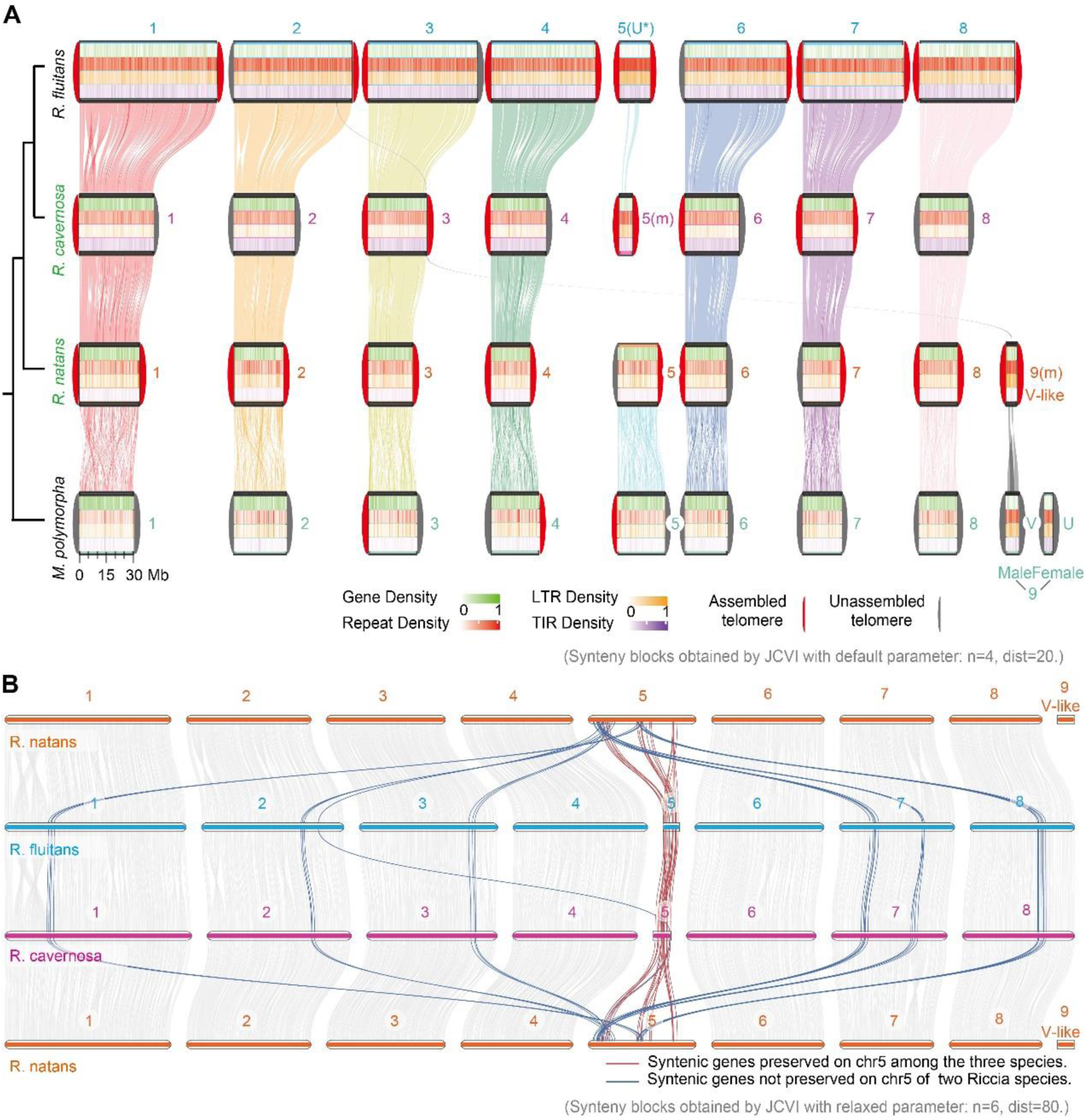
Chromosome evolution in Marchantiophyta. **A)** Comparison of genome sequences from the four liverwort species. Gap-free telomere-to-telomere level genomes of the *R. fluitans* female, and the monoicous *R. cavernosa* and *R. natans* were assembled in the present study, and the *M. polymorpha* chromosome assemblies were retrieved from https://marchantia.info/ (the public version: MpTak v6.1). Chromosomes with IDs are displayed as boxes, with sizes indicating the chromosome lengths, showing the small sizes of the *M. polymorpha* U and V. Gene density, repeat density, LTR density, and TIR density are indicated by green, red, orange, and purple heatmaps, respectively, and the linking lines indicate syntenic blocks. The *R. fluitans* chr5, which we infer is a U chromosome, or was so recently, is indicated by U*. Assembled and unassembled telomeres are indicated separately at each box end in red and gray, respectively. The links between each chromosome show syntenic blocks among them (based on protein-coding genes, using MCscan JCVI utility libraries with the default parameter values). **B)** Synteny analysis of just the Ricciaceae species, *R. fluitans*, *R. cavernosa*, and *R. natans*, to reveal syntenic blocks not distinguished by the analysis in Part A (parameter values dist=80 and n=6). Numbered horizontal bars represent chromosomes, and linking lines indicate syntenic blocks based on protein-coding genes with likely orthologues (defined as reciprocal best hits) in at least two of the three Ricciaceae species, using MCscan JCVI utility libraries, with the parameter values: dist=80, n=6, -full. Coloured lines indicate colinear blocks with at least three chr5 genes (red for intra-chromosomal rearrangements among the three species, and blue for inter-chromosomal ones in the most recent common ancestor of *R. fluitans* and *R. cavernosa*).

**Table 1.**
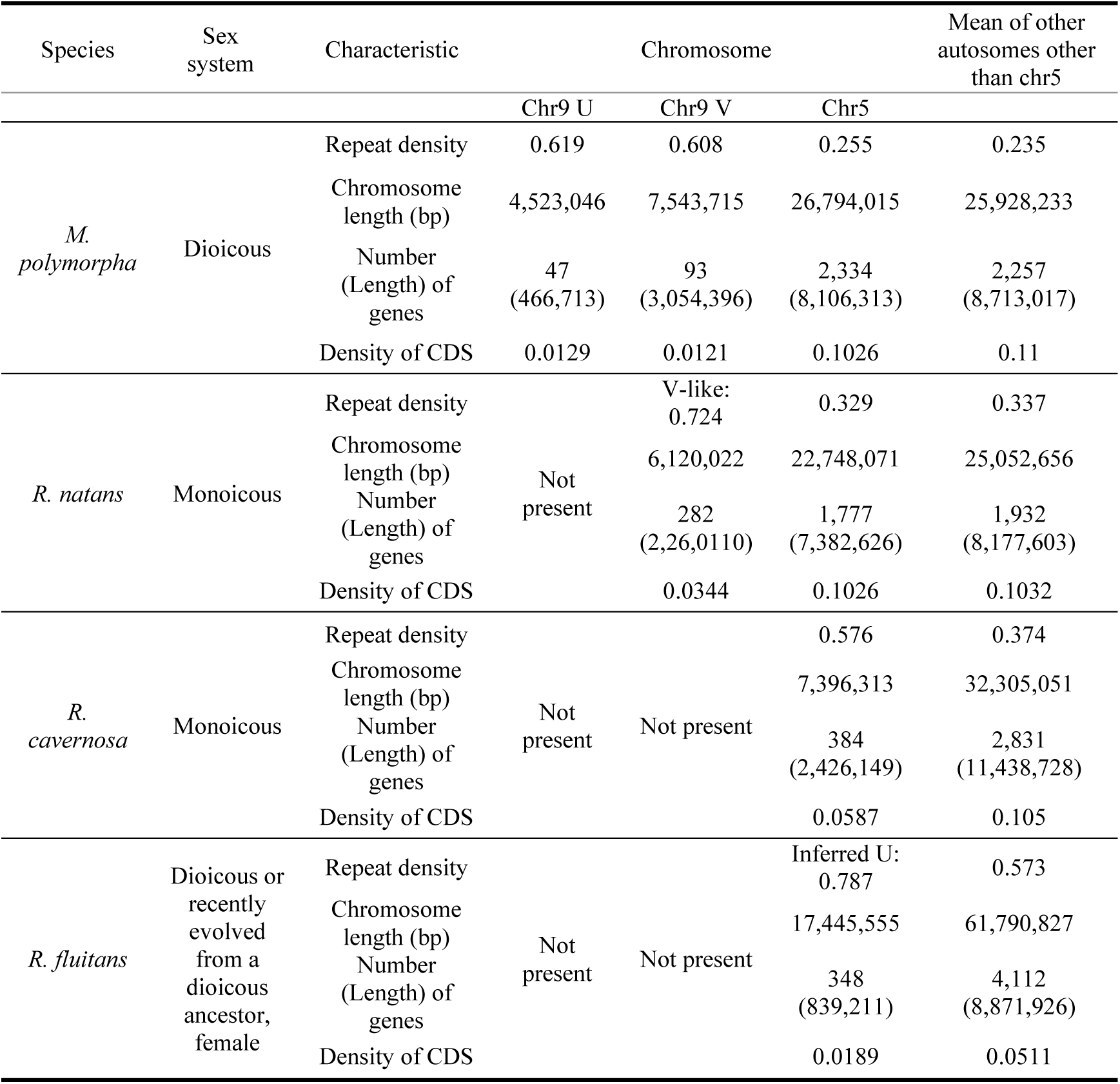
Comparisons of repetitive sequence contents of different liverwort chromosomes.

### Karyotype differences inferred from chromosome assemblies

Liverworts vary in chromosome number and some have micro-chromosomes (denoted by *m* in karyotype descriptions; Siler, 1934; Berrie, 1963; Smith, 1979; Althoff and Zachgo, 2020). In the species investigated here, the karyotypes are 8+m in *R. natans* (Siler, 1934), versus 7+m in *R. cavernosa* (Na-Thalang, 1980) and *R. fluitans* (Wyatt and Davison, 2013); in the right-hand part of Fig. 1, we classified the *m* chromosome of *R. fluitans* as a possible U, based on evidence from our genome sequences, as described below. These chromosome numbers suggest loss of one chromosome in the common ancestor of the two *Riccia* species. The observation of micro-chromosomes in both the monoicous species, *R. natans* and *R. cavernosa*, also suggests that each could be descended from a sex chromosome in a dioicous ancestor, possibly both from chromosome 9 (chr9), which is the sex chromosome in *M. polymorpha* (8+U/V sex chromosomes), and is a potentially ancestral liverwort U/V chromosome (reviewed in Singh et al., 2023).

Although *M. polymorpha* diverged from the Ricciaceae over 167 mya (Fig. 1; Fig. S4), the genome architectures are largely conserved between these distantly related liverworts. Synteny analysis nevertheless revealed extensively collinear homologous chromosomes in all four species (Fig. 2A), consistent with a recent comparison of *M. polymorpha* and *R. natans* (Singh et al., 2023). Intra-chromosomal rearrangements were detected in all three species compared with *M. polymorpha*, but no large inter-chromosomal ones. Both analyses (Singh et al., 2023 and our study), show that the *R. natans* micro-chromosome indeed carries homologous regions exclusively with the *M. polymorpha* V sex chromosome (Fig. 2A), supporting the idea just mentioned that both are descended from a common ancestral V. Since *R. natans* is monoicous, we describe this chromosome as V-like. The chr9 V-like chromosome is small (Table S3.3), similar to the small *M. polymorpha* V (Fig. 2A). Notably, neither study detected an intact U homologue in *R. natans*.

In contrast, searches using the same criteria as for *R. natans* (see Methods) detected no segments homologous to *R. natans* or *M. polymorpha* chr9 sequences in either of the two *Riccia* species, though collinear blocks were identified with all the other *M. polymorpha* chromosomes. This suggests that the V-like chr9, as well as the U, has been lost from the *Riccia* species.

Another striking genomic difference observed in *Riccia* is that chromosome 5 is very small in both species (Fig. 2A; Table S3.3). As the *Marchantia* sex chromosomes, and those of other haploid plants, are often remarkably small (Silva et al., 2021; Yu et al., 2022), this suggests that chr5 might represent a new sex chromosome that evolved in a turnover event, like those detected in many other organisms (Vicoso, 2019). More detailed synteny analysis with MCscan (JCVI utility libraries) of just the three non-*Marchantia* species (defining colinear blocks using the parameters in the Fig. 2 legend), confirmed the homology of the small *R. fluitans* chromosome and the micro-chromosome of *R. cavernosa* with the larger *R. natans* chr5 (Fig. 2B; the homologous blocks found in all these three Ricciaceae chr5 assemblies total about 10.7 Mb, or 61.49% of the *R. fluitans* chromosome’s length). Clearly, therefore, this micro-chromosome in *Riccia* is not the homologue of the micro-chromosomes of *M. polymorpha* (its chr9 V), or of the *R. natans* V-like chr9. Taken together, these findings are consistent with a transition from monoicy to dioicy in *Riccia*, after loss of the old sex chromosome, chr9, with possible evolution of a new sex micro-chromosome pair from the ancestral autosome chr5. We next describe other results consistent with this possibility, before evaluating it further.

Remarkably, given the rarity of inter-chromosomal rearrangements in liverworts (Linde et al., 2023), including the species studied here (Fig. 2A), blocks of sequence that are on chr5 in the closest outgroup, *R. natans* are scattered across five different autosomes in both *R. cavernosa* and *R. fluitans* (Fig. 2B); about 5.7Mb in total, corresponding to 25.17% of the *R. natans* chr5 length, is in a different location in *Riccia.* In both *Riccia* species, six of these segments are in similar locations on chr1, chr2, chr3, and chr8, and two are on chr7. All these changes (except for a second micro-translocation to chr2 in *R. fluitans*, after its divergence from *R. cavernosa*) must have occurred in a common ancestor of *R. fluitans* and *R. cavernosa*, before these species split, though their timing is not precisely known. These changes closely resemble the translocations inferred in the monoicous species *R. natans* (Singh et al., 2023), in which the chr9 U sex chromosome genes have moved to other chromosomes, and chr9 V sex chromosome has been retained (as a micro-chromosome). They are thus consistent with a period of monoicy in the *Riccia* lineage.

### Evidence that chromosome 5 is non-recombining in both *Riccia* species

To further evaluate whether chr5 could indeed be the U chromosome of *R. fluitans*, or of a dioicous *Riccia* ancestor, we analysed repetitive sequences, since high repetitive sequence density is evidence that a chromosome, or chromosome region, recombines rarely, or is completely non-recombining (Charlesworth et al., 1994; Charlesworth et al., 2005). Sex-linked regions and sex chromosomes (which do not recombine) therefore usually have higher repeat sequence densities than autosomes (and consequently lower gene densities than autosomes) (Kejnovsky et al., 2009; Ellegren, 2011; Zhou et al., 2020); consequently an unusually high repetitive sequence density of a chromosome would support the hypothesis that it is a sex chromosome. In both *Riccia* species Chr5 is indeed an outlier in this density, and its gene densities are lower than for the autosomes (Tab. 1).

Furthermore, chr5 carries unusually small absolute numbers of genes. In *R. fluitans*, it carries only 348 genes, many fewer than the other chromosomes, and the *R. cavernosa* micro-chromosome carries only 384. These properties resemble those of the (non-homologous) *R. natans* V-like chr9 (282 genes), and the *M. polymorpha* V chr9, which again differ strikingly from the other chromosomes in these characteristics (Tab. 1; Table S3.3).

Another predicted characteristic of sex chromosomes is low sequence diversity, a consequence of hitch-hiking processes in the absence of recombination (Hill and Robertson, 1966; Felsenstein, 1974), including the spread of advantageous mutations (selective sweeps) and/or elimination of deleterious mutations (Charlesworth et al., 2005; Kaiser and Charlesworth, 2010; Charlesworth, 2012). These effects have commonly been detected as lower nucleotide diversity, which shows the lower diversity values for sex chromosomes than autosomes (Qiu et al., 2010; Larracuente and Clark, 2014). In a haploid plant, such a diversity difference indicates that some sexual reproduction occurs (otherwise the whole genome would have a uniform low diversity), and this can be tested particularly clearly as the phase of sequence variants is known. *M. polymorpha* shows extensive vegetative reproduction (Sandler et al., 2023), as suggested for *R. fluitans*; nevertheless the comparison with the autosomes indicates that its sex chromosomes recombine especially infrequently (Fig. S6). If *R. fluitans* occasionally reproduces sexually, its females’ U chromosome should show a similarly low diversity. Resequencing of three *R. fluitans* accessions (from geographically distant locations) indeed yielded similar results for its chr5 to those for chr9 in *M. polymorpha*. The median nucleotide diversity was 0.005 across chr5, versus 0.0245 for the other chromosomes (Fig. S6). As just noted, this is evidence that some sexual reproduction occurs, consistent with the suggestion above that this species became clonal recently. Importantly, low diversity is found across the whole of chr5, showing that it does not simply reflect a pericentromeric location of a non-sex chromosome, where low diversity is consistently observed (Charlesworth, 2019); *R. fluitans*’ wholly or largely vegetative reproduction also makes introgression unlikely, arguing against the hypothesis that a genome region from a related species with a different arrangement spread, causing a high frequency of heterozygotes and a low recombination rate in the region. Low diversity is, however, expected for a fully sex-linked region.

All our findings therefore tend to support the hypothesis that chr5 became a new sex chromosome pair in a dioicous *Riccia* ancestor, and stopped recombining long enough ago to have evolved high repetitive content and small gene numbers, resembling the sex chromosomes of other bryophytes reviewed above.

The gene movements described above, from chr5 to other chromosomes within *Riccia*, reducing the size of chr5, resemble those from the ancestral sex chromosome chr9 to autosomes in *R. natans* (Singh et al., 2023). We suggest that the convergent evolution of small gene numbers (not entirely due to repeat-richness) in chr5 and chr9, reflect both having been sex chromosomes for part of their evolutionary history. Some of the gene losses could reflect movements off a former sex chromosome during periods of monoicy, as documented in *R. natans* (Bowman, 2016), but complete loss of chr9, creating the observed *Riccia* state with only 8 chromosomes plus chr5, would require loss of V-linked genes important for male functions, which was argued to be unlikely (Singh et al., 2023). The complete loss of the ancestral chr9 is therefore consistent with evolution of a new sex-determining system. Other genes losses would probably have occurred during the evolution of dioicy (similar to those that led to the small gene number of the *M. polymorpha* UV chr9 pair mentioned above); such changes are not observed for autosomes in these or other species. Our observations are thus consistent with dioicy having re-evolved in a *Riccia* ancestor. If *Riccia* had remained monoicous, and chr5 had remained a non sex chromosome, its sex chromosome-like properties described above remain inexplicable.

### Searches for genes from the UV pair prior to the evolution of *Riccia*

If chr5 is, or has been, a sex chromosome in *Riccia*, it is interesting to ask whether it has gained sex-determining genes from an ancestral chr9 homolog. In *M. polymorpha*, Iwasaki *et al*. (Iwasaki et al., 2021) annotated 23 and 39 genes, respectively, with high confidence as U- or V-specific, defined as genes without autosomal paralogs that might reflect gene movements from or onto a sex chromosome. We reanalyzed these genes using blast searches for complete protein-coding sequences, followed by phylogenetic analysis, identified autosomal paralogues for 4 U and 3 V genes in *M. polymorpha* (Tables S5.1 and S5.2; Fig. S7, which used Carey et al., 2021b’s high-quality genome assembly of *Ceratodon purpureus* as an outgroup to further validate inferences of autosomal paralogues in *M. polymorpha*, and of their orthologues in other Marchantiales species).

Most *M. polymorpha* U-specific genes have clearly been lost in the Ricciaceae, consistent with the loss of the entire ancestral U chromosome in *R. natans* (Bowman, 2016). Only five genes (5/19 = 26.32%), MpUg00040, MpUg00110, MpUg00330, MpUg00290, and MpUg00370 (Mp*BPCU*), have orthologues identified in at least one of the three Ricciaceae species investigated (Fig. 3E; Table S5.1), and only the MpUg00110 orthologue is on chr5 in *R. fluitan*s; those of the other genes are on various autosomes (Fig. 3E; Table S5.1). *R. natans* has all five of these genes, whilst two and four of them are found in *R. cavernosa* and *R. fluitans*, respectively (Figs. S8 and S9; Tables S5.1, S5.4 and S5.6). We confirmed the finding of Singh *et al*. (2023) that two of these genes (the MpUg00110 and MpUg00330 orthologues) are close together on chr4 in *R. natans*, and found that another, MpUg00040, is on *R. fluitans* chr4 (Fig. 3E, Fig. 10A; Table S5.5). A fragment of the ancestral U chromosome may thus have translocated to the autosomal chr4 in the common ancestor of Ricciaceae and then either translocated to a region near the telomere of chr4 (its present location in *R. natans*) or become lost (Fig. 10A; Tables S5.3, S5.7 and S5.8).

**Figure 3.**
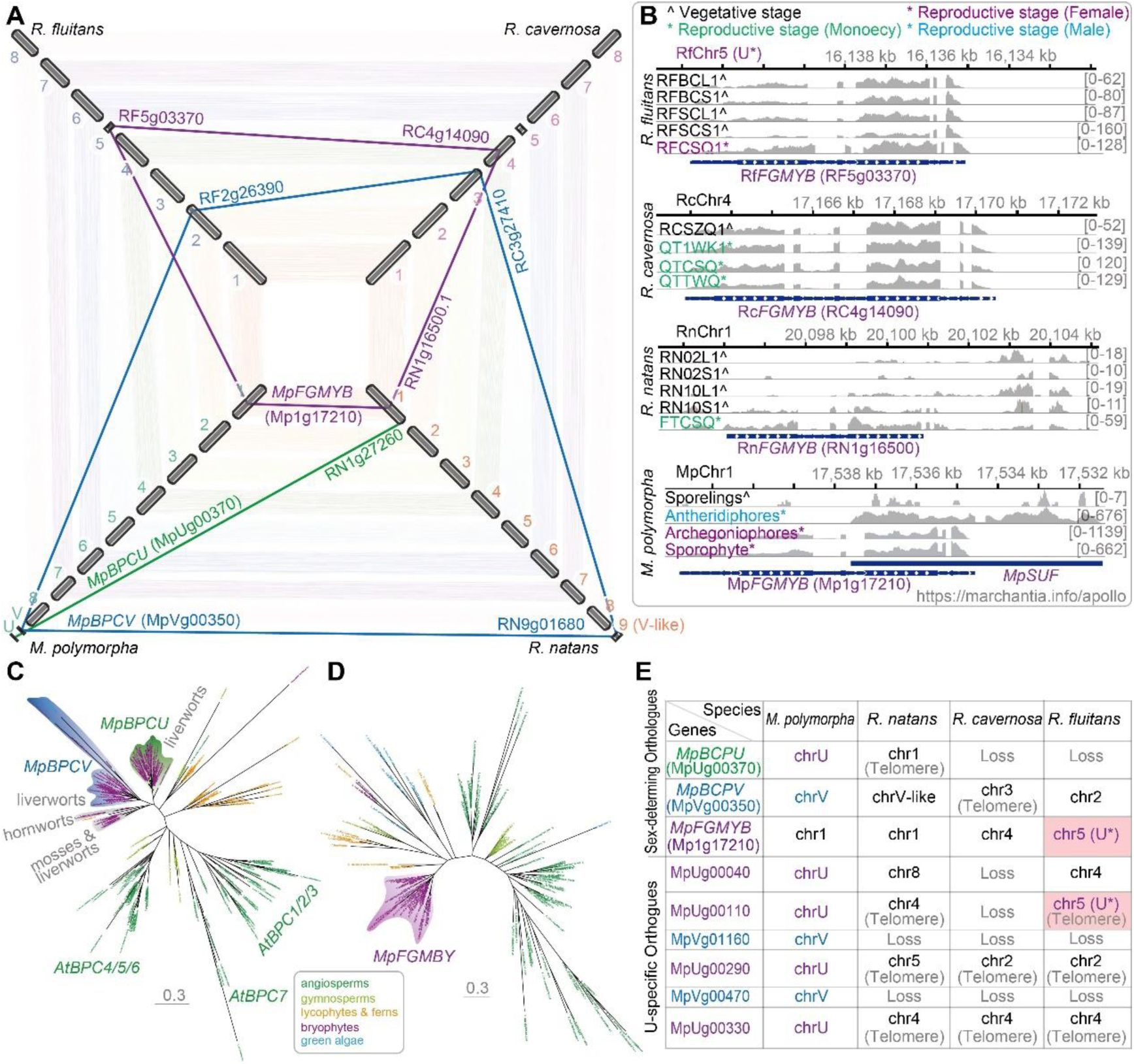
Evolution of genes known to be associated with sexual differentiation in *M. polymorpha*. **A)** Identification of orthologues of sexual differentiation-associated genes in the chromosomes of the four liverwort species studied here. The wide coloured bars represent chromosomes with their numbers indicated, and the positions of the orthologue of each of the 3 genes known to be involved in the sex-determination pathway in *M. polymorpha* are indicated by thin lines joining the copies in the three different Ricciaceae species, with the gene names for each species indicated at their two ends; the genes are also colour-coded as follows: *FGMYB* (blue), *BPCV* (green), and *BPCU* (cyan). **B)** Expression of *FGMYB* and of the *SUF* orthologues in different tissue samples from *M. polymorpha* (https://marchantia.info/apollo). Blue bars indicate the organization of the *FGMYB* and *SUF* genes in *M. polymorpha*, and blue bars with white arrowheads represent coding regions. Expression in the Ricciaceae species is also shown, at stages indicated by the colour coding at the top (Table S2.5 provides details of the samples). Blast searches of the whole genome for *SUF* orthologues detected a similar sequence only in *M. polymorpha*, and expression analysis found no RNA-seq reads mapping to the *SUF* locus in any of the three Ricciaceae species. Numbers at the right of each row in the diagrams for each species show the total numbers of reads. **C)** Phylogenetic analysis of *BPC* genes in green plants. The *BPCU* clade is found only in liverworts, while genes in the *BPCV* clade are present in liverworts and algae. **D)** Phylogenetic analysis of *FGMYB* genes in green plants. **E)** Summary of the fates of genes associated with sexual differentiation. The possible *R. fluitans* U chromosome is indicated by U*.

A U chromosome can potentially be lost after female-essential U-linked genes become duplicated to autosomes, or if different autosomal genes can perform their functions (provided that this does not have major detrimental effects on male sex functions). This appears to have occurred in *R. natans*, with loss of the chr9 U. A plausible hypothesis, suggested for the loss of the U in *R. natans* (Singh et al., 2023) is that the evolution of monoicy involved an initial genotype carrying both a U and a V, which selected for loss of one of the chromosomes.

In contrast, most *M. polymorpha* V-specific genes (29/36 = 80.56%; Tables S5.2, S5.9 and S5.10) have been retained in all three Ricciaceae species studied here (Figs. S9, S10B and S11; Tables S5.2, S5.7 to S5.11). *R. natans* has 28 orthologues, 25 of them still on the single V-like chromosome, and only 3 on autosomes. The total numbers of such genes are similar in *R. cavernosa* (27) and *R. fluitans* (25), but none of them are on chr5, the *R. cavernosa* micro-chromosome, or the possible *R. fluitans* U (Table S5.10). V chromosome genes in *M. polymorpha* and other bryophytes are likely to have important male gametophyte functions, such as sperm motility functions, which will prevent their loss, as monoicous species also require these functions for sexual reproduction, including self-fertilisation (Bowman, 2016). Likely male function-essential genes are distributed across the V chromosome, and deletion of large V-linked regions would therefore be disadvantageous, explaining their retention in *R. natans* (Singh et al., 2023).

Since the lineage that includes *R. natans* split from the *Riccia* species before the most recent ancestor of *R. cavernosa* and *R. fluitans*, it is likely that these species shared an ancestral V-like chromosome that has nevertheless become lost in *Riccia*. To determine the fate of this chromosome in *Riccia*, we searched for genomic fragments homologous to the *R. natans* V-like chr9 by inspecting the genomic locations of inferred orthologues of *M. polymorpha* V sex chromosome-specific genes in all three Ricciaceae species. Two homologous autosomal segments were detected in *R. cavernosa* and *R. fluitans*. One region, near the telomeres of chr6 in both *Riccia* species, includes four such genes (Fig. S10B). This suggests translocations of chr9 genes to this autosome in the common ancestor of *R. cavernosa* and *R. fluitans*, probably before dioicy re-evolved within the *Riccia* lineage. Eight other putative V gene orthologues were found. In *R. cavernosa*, they are in a genomic region close to the chr3 telomere, whilst in *R. fluitans*, five are on chr2 (Fig. S10B; Table S5.11). These results suggest further movement that occurred independently after the split of these two species.

### Rearrangements of chromosome 5 in *Riccia*

A further property of the *R. fluitans* and *R. cavernosa* chr5 that is convergent with the sex chromosomes of many taxa (including the bryophytes *M. polymorpha* and *C. purpureus*) is that, unlike the other chromosomes, gene order is considerably rearranged, in this case, between the two species (like the low sequence diversity described above, this affects the whole chromosome; Fig. S10C; Table S3.4). If the *R. fluitans* chr5 represents a *Riccia* U-like chromosome, the *R. cavernosa* chr5 could therefore represent a *Riccia* V-like chromosome, with each of these lineages having inherited a different sex chromosome from a common dioicous ancestor with chr5 as its UV pair. This would resemble the situation in the genus *Volvox* (a haploid plant lineage distantly related to bryophytes), in which one functionally monoicous (homothallic) lineage carries only a U and another carries only a V chromosome (Yamamoto et al., 2023). This suggested that homothally of *Volvox africanus* was probably initiated by a stage with both a U and a V haplotype present, as also suggested for the evolution of monoicy in *R. natans* (Bowman, 2016).

We tested this possibility further by estimating divergence between coding sequences in the two *Riccia* species. Under this scenario, U and V genes should have diverged in the dioicous ancestor before genes on the other chromosomes started diverging (which would have started only when the species split). The species diverged too long ago to use synonymous sites (Ks) to test this, as substitutions are saturated (the mean Ks value for every chromosome is >50%, Table S6). We therefore estimated divergence per non-synonymous site (Ka). The mean for chromosomes other than chr5 is 8.73%, and the chr5 value is 23.32% higher (Table S6), consistent with descent from the members of a UV pair whose sequences had their last common ancestor in a common ancestor of these species. We cannot exclude the alternative that the set of genes on this chromosome happen (for unknown reasons) to evolve faster than those on other chromosomes; however, this seems unlikely, as 46 syntenic genes were detected and analysed on chr5 (Table S6). Further arguing against this alternative, divergence estimates for chr5 homologs between *R. natans* and *M. polymorpha* are lower for chr5 than for other chromosomes. Therefore, chr5 sequences are not simply fast-evolving (though many of them are no longer found on the small *Riccia* chr5, so that the two inter-species comparisons are not directly comparable).

### Evolution of genes involved in sex determination

On the hypothesis that a new sex chromosome evolved in a *Riccia* ancestor, *Riccia* might either have evolved new sex-determining genes, different from those in *M. polymorpha*, or the *M. polymorpha* sex-determining system might have been retained, with the genes moving onto chr5. Presence of a dominant U chromosome locus controlling female versus male sex specification in the development of gametophytes, termed feminizer, has been indicated in phylogenetically distant liverworts, including *Marchantia*, *Pellia*, and *Sphaerocarpos* (Haupt, 1933; Lorbeer, 1936; Heitz, 1949; Singh et al., 2023). In *M. polymorpha*, this genetic sex-determining function involves the U chromosome gene MpUg00370, which encodes a transcription factor BASIC PENTACYSTEINE ON THE U CHROMOSOME. The feminizer is therefore also referred to as Mp*BPCU* (Iwasaki et al., 2021). Mp*BPCU* expression in female gametophytes regulates an autosomal downstream locus encoding a key regulator, *FEMALE GAMETOPHYTE MYB* (Mp*FGMYB*), essential for female sex organ formation. Expression of Mp*FGMYB* is suppressed in *M. polymorpha* males by a *cis*- acting antisense long non-coding RNA, *SUPPRESSOR OF FEMINIZATION* (Mp*SUF*), from the same locus (Hisanaga et al., 2019). In female gametophytes, the repression is removed by Mp*BPCU*-mediated suppression of Mp*SUF* transcription, leading to development of female sex organs.

We investigated the evolutionary fates of these genes in the Marchantiales species studied here. The orthologue of the feminizer *BPCU* is retained in the monoicous *R. natans*, but it is on chr1, an autosome (Fig. 3A). In both of the later-branching *Riccia* species, the *BPCU* and *SUF* orthologues are both absent (Figs. 3A to 3C; Fig. S12), suggesting that a different system, not involving feminizer and suppressor, evolved to determine sex in their inferred dioicous ancestor. However, as might be expected, orthologues of the downstream-acting *FGMYB* gene were detected in all the genomes investigated (Figs. 3A and 3D; Fig. 13), and *cis*-acting transcriptional control of *FGMYB* by the *SUF* gene may still occur in the monoicous *R. natans* (Fig. 3B). In *M. polymorpha*, the *SUF* and *FGMYB* genes partially overlap, but their transcripts can be distinguished using non-overlapping regions. However, in *R. cavernosa* and *R. fluitans*, no reads mapping to the *FGMYB* flanking regions were detected on either strand, suggesting absence of any functional *SUF* gene in these two species. Intriguingly, in *R. fluitans*, the *FGMYB* gene, which in *M. polymorpha* is autosomal, is on chromosome 5, as is a gene orthologous to the *M. polymorpha* MpUg00110 gametolog, suggesting possible movements of these genes onto a new sex chromosome. Taken together, these observations suggest that a new and as yet unidentified master sex-determining gene evolved to establish genetic control of sexual dimorphism in *R. fluitans*, or its dioicous ancestor.

## Discussion and Conclusions

Overall, the results of our analysis suggest that sex chromosomes and sex determination may not be conserved in all liverworts, as *Riccia* may have evolved a new sex chromosome, homologous with chr5 in other lineages. Our study of *Riccia* also uncovered unexpected evolutionary changes affecting U and V sex chromosomes, including movements of genes to the autosomes. The reason(s) for the appearance of formerly sex-linked genes in autosomal locations are not yet fully understood. First, as noted above, the U and V chromosomes in dioicous species are often diminutive, and often denoted by “*m*-chromosomes” (for miniature or micro). Our genome sequences reveal the likely evolution of a diminutive U chromosome 5 paralleling the evolution of the diminutive UV chr9 pair in *M. polymorpha*.

Suggestions that some bryophyte *m*-chromosomes might be sex chromosomes (e.g. Allen, 1945) have not previously been tested, because, without genome sequencing, it is not possible to show that these chromosomes carry homologs of genes on defined chromosomes of related species. These might therefore be supernumerary elements such as B chromosomes, especially as both these chromosome types are heteropycnotic and heterochromatic (Anderson, 1984). The relationship between the micro chromosome 9 UV pair of *M. polymorpha* and the V-like one in *R. natans* is now clear (Singh et al., 2023), and our findings suggest that the *Riccia* chr5 represents a similar situation, although no definitively dioicous species with a chr5 UV pair has yet been studied, this state may have existed only in an ancestral *Riccia* species. Our suggestion is based both on the properties the *Riccia* chr5 shares with other bryophyte sex chromosomes, including a complete lack of recombination (for which the previous section describes evidence, and which can explain their repeat-rich and heterochromatic nature), and the convergent extraordinarily small number of genes, compared with other chromosomes. Both sex chromosomes and accessory chromosomes may share this property, but for different reasons, as the case of the *Riccia* chr5 illustrates. Accessory chromosomes are generally made up of a mosaic of sequences from all, or most, of the “A chromosome” set (Makunin et al., 2018; Blavet et al., 2021). In contrast, the *R. fluitans* chr5 gene content does not consist of homologs from other *R. fluitans* chromosomes (Figure 2B). Instead, it shares homologous regions with chr5 of *R. cavernosa*, indicating that this chromosome was inherited by these species from a common *Riccia* ancestor; the outgroup, *R. natans*, allows us to infer that parts of this chromosome moved from chr5 in *Riccia*, rather than onto it, as occurs for B chromosomes, which incessantly gain genes from the A chromosomes. The change to being a sex chromosome is further supported by the observation that chr9, which is the sex chromosome in the outgroup species, is missing from the *Riccia* genome, and our analyses of gene content show that this is not due to a fusion with an autosome.

Gene losses from newly evolved V and U chromosomes, causing reduced physical sizes, are not unexpected. First, they are a predicted consequence of a lack of recombination. In the case of V and U chromosomes in haploid plants, genes essential for male or female sexual differentiation functions, respectively, must be retained (Bull, 1978), but genes required only by sporophytes can be lost from the U or the V during sex chromosome evolution (though Bull predicted that neither sex chromosome should show major degeneration). Second, as described above, gene movements have clearly also occurred after monoicy evolved. The reduction of the chromosome number from 9 to 8 in *Riccia*, involving eventual complete loss of the chr9 V, with its genes being on other chromosomes, may reflect independent evolution under monoicy in this lineage, as well as in the *R. natans* lineage (see Fig. 4). Gene losses from the U or V, in both the evolution of dioicy, or its breakdown to monoicy, may sometimes involve duplications to autosomes, but such evolutionary changes have rarely been studied until now.

**Figure 4.**
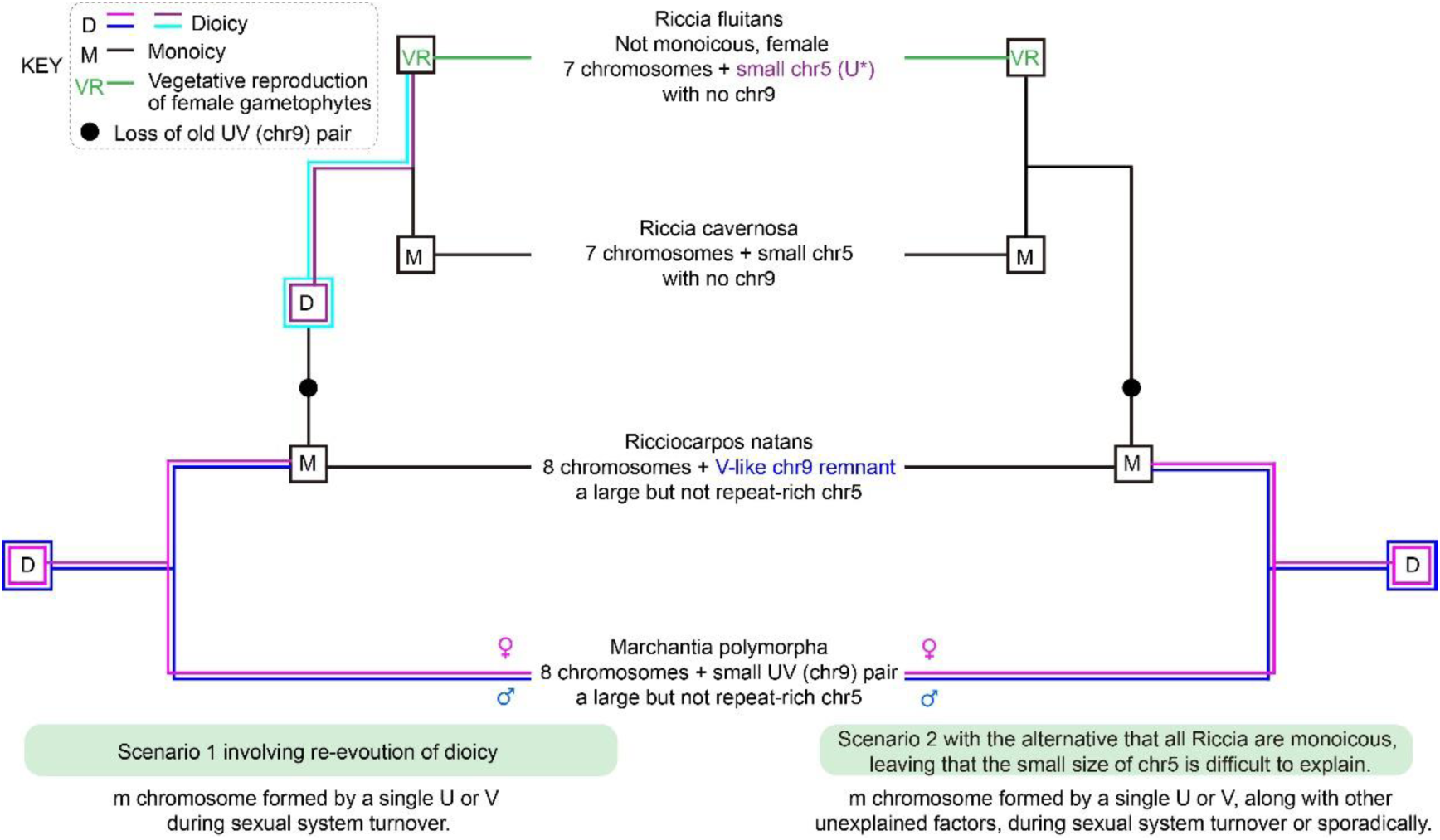
Evolutionary changes affecting the sex chromosomes during transitions between dioicy and monoicy in liverworts. The inferred sexual systems of the species studied are indicated at different important time-points by the letters shown in the key. U* indicates the *R. fluitans* possible U chromosome. A pair of lines in the phylogeny indicates U and V sequences in dioicous species, and a single line indicates monoicy (or loss of a U or V chromosome). For dioicous lineages, pink and blue indicate a chr9 UV pair, while red and turquoise indicate the proposed chr5 UV pair. The dioicous *M. polymorpha* has 8 chromosomes plus a small highly diverged UV pair, consistent with the evolution of sex chromosomes in bryophytes involving chromosome size diminution. The monoicous *R. natans* karyotype is similar (in both these species, chr5 is large, and not repeat-rich), but *R. natans* has no U (some ancestral U genes are autosomal, as explained in the text) and has only a V-like chr9 remnant. This suggests that evolution of monoicy can involve further diminution, and even loss, of sex chromosomes, making it difficult to separate the alternative causes of small chromosomes outlined in the text. The left-hand part shows the possibility that, after the complete loss of chr9 (which is absent from both *Riccia* species), dioicy re-evolved at an unknown time. This created a new UV pair with the sex chromosome-like properties of loss of recombination (indicated by its high repetitive content and low sequence diversity), and diminutive size, involving regions moving to various autosomes. Alternatively (as in the right-hand diagram), dioicy did not evolve in *Riccia*, but monoicy was maintained (and females in *R. fluitans* evolved by loss of male function, which is potentially possible, though unlikely, as explained in the Introduction section of the text). The loss of chr9 in *Riccia* can potentially be explained under this continued monoicy scenario, which might have led to further loss of chr9 genes that are still present in the *R. natans* V-like chr9 remnant. This would require loss of V-linked genes important for male functions, and thus to the state with only 8 chromosomes plus chr5. However, the changes in chr5 are inexplicable unless it was a sex chromosome.

Independent dioicy to monoicy transitions in these haploid plants suggest that (just as in diploid plants) the presence of separate sexes in a set of lineages need not imply that dioicy evolved in a common ancestor and remained under the same control in all extant species. As mentioned above, transitions from ancestral dioicy to monoicy are likely during the evolution of *Riccia*. Fig. 4 summaries two possibilities for this genus. Scenario I is the possible re-evolution of dioicy, followed by evolution of small sex chromosomes (carrying few genes, often with sex-specific functions, partly due to translocations of genes to autosomes), and rearrangements after recombination stopped. Under the alternative Scenario, II, dioicy did not re-evolve in *Riccia*, and chr5 did not become a sex chromosome; instead, females in *R. fluitans* evolved by loss of male functions in a monoicous ancestor. The changes in chr5 then remain unexplained and mysterious. It seems unsatisfactory to suggest that, for unknown reasons, the chr5 coincidentally stopped recombining, gained a high repetitive content, and started to lose genes and became a micro-chromosome. Further analysis of more species from *Riccia* are needed to definitively distinguish between these possibilities, but our overall interpretation that dioicy probably re-evolved in an ancestor of the two *Riccia* species, with *de novo* evolution of a new sex chromosome pair, is strongly suggested based on the striking and unusual evolutionary properties of the *R. fluitans* chr5 which are convergent with those of other sex chromosomes, including accumulation of repetitive sequences, lower sequence diversity and more rearrangements than other chromosomes, all of which indicate that recombination is restricted or entirely suppressed. A greatly reduced number of genes carried on the chromosome also parallels the predicted and widely observed genetic degeneration after a sex-linked region stops recombining. If confirmed, Scenario I implies that a new system controlling male and female gametophyte development may have evolved in *Riccia*, as the failure of our searches to detect several key *Marchantia* sex-determining genes in *Riccia* is consistent with this having happened.

## Methods

### Plant material collection and culture

The liverworts *Ricciocarpos natans* and *Riccia fluitans* were collected from Hangzhou Botanical Garden, Zhejiang, China. *Riccia cavernosa* was collected from a crop field of a farm at Beijing, China. Sterilized thalli were obtained by chemical disinfection of tissue surfaces using 0.4% sodium hypochlorite solution, followed by four washes with deionized H_2_O. The thalli were then cultivated in solid Knop medium plates (Schween et al., 2003) at 22 °C under long-day conditions (16 h light/8h dark). Then, isolates were established by proliferating tissues from single colony. Isolations of the three liverworts species used for our analyses were cultured on solid and liquid media of 1/2 Gamborg’s B5 (GB5) medium and 1/10 GB5.

### DNA and RNA extractions

High-quality genomic DNA was extracted from young thalli using cetyl-trimethylammonium bromide (CTAB) method (Gawel and Jarret, 1991). Tissues at various developmental stages were harvested for expression analysis and genome annotation. The total RNA was extracted using a RNeasy Plant Kit (Tiangen, Beijing).

### Genome size estimation

Genome sizes of *R. natans* and *R. fluitans* were estimated by flow cytometry to be 292.71Mb and 485.48Mb, respectively, using fresh thalli. Genome sizes of *R. natans* and *R. cavernosa* were estimated by k-mer analysis to be 198.00Mb and 240.85 Mb, respectively. Genomic DNA samples were sequenced by a combined strategy as follows. A total of 32Gb (∼156-fold coverage) and 62Gb (∼265-fold coverage) short reads were respectively generated by BGI-seq T7 or Illumina Hi-seq 2000 platforms for *R. natans* and *R. cavernosa*. Reads with low quality were filtered by Trimmomatic v0.39 (Bolger et al., 2014) (parameter: “LEADING:20 TRAILING:20 SLIDINGWINDOW:4:20 -threads 48 MINLEN:36”), and quality was checked by FastQC v0.11.9 (Wingett and Andrews, 2018). Resultant clean reads were used to estimate genome size by *k*-mer using KmerGenie v1.7051 (Chikhi and Medvedev, 2013) with default parameter.

### Genome assembly

Circular consensus sequence (CCS) HiFi long reads for use in assemblies were produced by PacBio with a 15-20kb insert library (CCS reads N50: ∼17kb). A total of 36Gb (∼176-fold coverage) of PacBio reads were obtained for *R. natans*, 30Gb (∼66- fold coverage) for *R. fluitans*, and 36.5Gb (∼156-fold coverage) for *R. cavernosa*, and were used for *de novo* assemblies with Hifiasm software v0.16.1-r375 (Cheng et al., 2022) with the default parameters. To identify contaminant sequences, we downloaded 405 fungal genomes, 24,189 bacterial genomes, 496 archaeal genomes, and 11,664 viral genomes from NCBI. We then constructed nucleotide databases for each group and aligned the raw assembly results to these four databases (using BLASTN (Nucleotide-Nucleotide BLAST 2.12.0+) (Camacho et al., 2009) with an e-value ≤ 1e-5) to remove contaminant sequences. To remove organellar sequences from the genome, we used the same method, but with databases using the mitochondria and chloroplasts of *Marchantia paleacea*. Finally, aligning the remaining short contigs to the longer contigs showed that all of the former were repetitive fragments of long contigs, which we removed. The final *R. natans*, *R. fluitans* and *R. cavernosa* assemblies were composed of 12, 9, and 14 contigs, respectively. We also employed further parameter sets in Hifiasm and the Canu software v2.2 (Nurk et al., 2020) to assemble more versions of genomes for use in the gap filling step.

To obtain chromosomal assemblies, we used Hi-C assisted assembly technology, a development of chromosome conformation capture (3C). Fresh plant samples were treated with 1-3% formaldehyde at room temperature for 10-30 minutes to crosslink chromatin. The genome was then digested using the restriction enzyme *Dpn* II. The Hi-C libraries was prepared following the standard protocol for Hi-C library construction (Xie et al., 2015), and sequenced on an Illumina Hiseq 2000 platform (PE150), generating 35Gb (∼170-fold coverage) of *R. natan* sequences, and 67Gb (∼149-fold coverage) for *R. fluitans*. Low-quality reads were removed with the HiC-Pro pipeline (https://github.com/nservant/HiC-Pro) and mapped onto the draft assembly contigs with SAMTools v1.6 (Danecek et al., 2021) and BWA v0.7.17- r1188 (Li and Durbin, 2009). Finally, the Juicebox pipeline (Durand et al., 2016; Dudchenko et al., 2017) was applied to further cluster the contigs into pseudochromosomes.

We used three methods to fill the 3, 1, and 6 gaps remained in the assemblies of *R. natans*, *R. fluitans* and *R. cavernosa*, respectively, following (Song et al., 2021). 1) We constructed contigs with Canu v2.2 (Nurk et al., 2020) or Hifiasm software, using HiFi reads with various parameters, and mapping them to the pseudochromosomes with minimap2 to identify overlapping regions and fill the gaps. 2) We mapped low-coverage PacBio subreads to the pseudochromosomes with Minimap2 v2.24-r1122 (Li, 2018) to identify overlaps and fill the gaps. 3) In highly repetitive regions where the previous methods were ineffective, we aligned HiFi reads to the pseudochromosomes using minimap2 and manually adjusted the breakpoints and repeat copy numbers, in 30-50 rounds of iterative adjustments, proofreading, and assessment. Minor errors in the copy numbers of repetitive sequences probably remain, but probably reflect individual variation and affect small proportions of the assemblies.

We assessed the completeness of the genome assemblies using BUSCO v5.4.3 (Seppey et al., 2019; Manni et al., 2021) and LAI (implemented in LTR_retriever v2.9.0; Ou et al., 2018).

### Assembly of organelle and bacterium genomes

After identifying the organellar and contaminant sequences in the previous steps, we obtained both circular and fragmentary genomes of these sequences. The organellar genomes were manually checked and reassembled based on the published chloroplast and mitochondrial genomes of *Marchantia paleacea*, downloaded from NCBI and aligned using minimap2 v2.24-r1122. The assembled chloroplast and mitochondrial genomes were annotated using CPGAVAS2 ( http://47.96.249.172:16019/analyzer/home; Shi et al., 2019).

Contaminant genomes were identified via BLASTN (Nucleotide-Nucleotide BLAST 2.12.0+; Camacho et al., 2009) aligning on NCBI contaminant database, as described above. This identifed three circular contaminant genomes from the *R. fluitans* dataset. Two of these were from *Mycobacterium intracellulare* and *Mycolicibacter minnesotensis*, and the third was from an unknown bacterium. A circular fungal contaminant genome was identified from the *R. natans* dataset, but we were unable to determine its species.

### Telomere structure

We searched for telomere sequences (TTACCC) within each genome using a local Perl script, and merged the sequences with BEDTools v2.30.0 (Quinlan and Hall, 2010) using the paremeters mergeBed -d 50; length ≥ 200bp (Table S4.2). All three *de novo* genome assemblies reached the gap-free and T2T chromosome level, and are nearly complete genomes.

### Transcript assembly

RNA-seq reads were generated by BGI-seq T7 platform, and low-quality sequences were processed and removed using Trimmomatic v0.39. The remaining cleaned reads were *de novo* assembled to transcriptome contigs via Trinity v2.8.5 (Grabherr et al., 2011). The full-length Iso-seq reads were generated by the PicBio CCS HIFI platform, which were used to collect the high-quality transcripts via IsoSeq3 (https://github.com/ylipacbio/IsoSeq3) using the refine and cluster models. The GMAP v20211217 (Wu and Watanabe, 2005) was used to map the full-length transcripts to the genome assemblies with default parameters. The StringTie2 v2.2.1 (Pertea et al., 2015; Shumate et al., 2022) and TransDecoder v5.50 (https://github.com/TransDecoder/TransDecoder) were together executed to assemble and annotate the transcriptomes using the short reads and long reads. Finally, gffread (https://github.com/gpertea/gffread) was used to identify the coding and protein sequences from the genome assemblies.

### Repeat annotation

Long terminal repeat (LTR) retrotransposons and miniature inverted transposable elements (MITE) were identified from the genome assemblies using the EDTA software (Extensive *De Novo* TE Annotator, v2.0.0; Su et al., 2021) with the default parameters. All repeat sequences identified were searched against the Swiss-Prot database (https://www.uniprot.org/) using BLASTX (Nucleotide-Nucleotide BLAST 2.12.0+) with 1E-10 to exclude matching non-TE proteins. We used BEDTools v2.30.0 to finding the location of repeats, and for a window of 100kb to estimate the repeat densities.

Simple repeats were masked by Ns and complex repeats were masked as lowercase in a genome.masked file to generate the softmask genome sequences using the same package (Campbell et al., 2014).

### Gene model annotation

MAKER-P MAKER v3.01.03 (Cantarel et al., 2008; Campbell et al., 2014) and PASApipeline v2.5.3 (https://github.com/PASApipeline/PASApipeline) were used to generate gene model annotations. As evidence for protein-coding genes, we used unique proteins encoded by the longest isoforms of each gene from five published genomes, including *Anthoceros angustus* (Zhang et al., 2020), *Arabidopsis thaliana* (Lamesch et al., 2011), *Ceratodon purpureus* (Carey et al., 2021), *Marchantia polymorpha* MpRak_v6.1r2 (Montgomery et al., 2020), and *Physcomitrium patens* v3.3 (Lang et al., 2018). The inchworms generated by Trinity v2.8.5 were used as expressed sequence tag (EST) evidence. Gene modelling was trained using Augustus v3.5.0 (Stanke et al., 2006) with the transcript sequences generated by Trinity v2.8.5 (Grabherr et al., 2011), and using GeneMark-ES-suite v4.* (Lomsadze et al., 2005) with the assembled genome sequences. To identify gene structures, all transcript sequences and gene models were integrated using the MAKER-P pipeline. The first round of MAKER-P output was subjected to SNAP (ZOE library version 2013-02-16 (Korf, 2004) gene model training, which was used in a second round of MAKER-P to refine the gene structures.

Annotations of spliced transcripts (isoforms) and untranslated regions (UTRs) in the whole genome sequences were based on the PASApipeline v2.5.3 (Haas et al., 2003; Campbell et al., 2006), incorporating the second round of MAKER-P output along with transcripts assembled from the iso-seq and RNA-seq reads.

### Annotation of gene function

Two methods were used to infer gene functions. The local version of eggNOG-mapper v2.1.9 (Cantalapiedra et al., 2021) was used to *de novo* predict the gene functional annotations based on precomputed orthology assignments. Then, BLASTP (Nucleotide-Nucleotide BLAST 2.12.0+) was used to align the proteins to *M. polymorpha* (https://marchantia.info/) and *A. thaliana* (https://phytozome-next.jgi.doe.gov/) gene sequences with an expectation value (*E*-value) of 1E-5.

### Evolutionary and Phylogenetic analyses

The three new genome assemblies generated in this study (*R. natans*, *R. fluitans*, and *R. cavernosa*), together with the published *M. polymorpha* one, and 53 published transcriptomic data sets (Dong et al., 2022) from all major liverwort lineages were used to infer the relationships in Fig. 1. Additionally, a total of 56 genome assemblies were used to infer the evolutionary history of green plants (Fig. 5; Table S4.1), mostly using genome and annotation data downloaded from Phytozome 13 (Goodstein et al., 2012), NCBI (https://www.ncbi.nlm.nih.gov/), and CNSA (Guo et al., 2020) with the latest version. Both phylogenetic analyses employed the same approach, as follows. We identified low copy-number orthologous genes using OrthoFinder v2.5.5 (Emms and Kelly, 2019) with the ‘-M msa’ option. These sequences were aligned by MUSCLE v5.1.linux64 (Madeira et al., 2022) and poorly aligned sequences were removed by TrimmAl v1.4.rev22 (Capella-Gutiérrez et al., 2009) with default parameters. The trees were constructed using IQ-TREE 2 v2.2.0.3 (Nguyen et al., 2014) to define conserved sites within these sequences, with automatic model selection and 1,000 bootstrap replicates.

To infer the divergence times between the four focal liverwort species, synonymous site divergence, *Ks*, was estimated between protein-coding genes identified in their genome sequences, using KSPlotting (https://github.com/EndymionCooper/KSPlotting). The divergence time was calibrated using the ages estimated (119.0-215.2 mya) between the genera of *Marchantia* and *Ricciocarpos* from Timetree (http://timetree.org/).

### Synteny analysis

Syntenic protein-coding gene pairs among the chromosomes of *R. natans*, *R. fluitans*, *R. cavernosa*, and *M. polymorpha* were identified using the MCscan python package of JCVI v1.2.7 (Tang et al., 2008) with default parameter values (“-n 4 --dist=20”; see Fig. 2A). Since no syntenic links found between chromosome 5s and other autosomes of three Ricciaceae species, more relaxed parameter were applied (“-n 6 --dist=80”; see Fig. 2B). Syntenic genes were identified using the parameters dist=80 to define windows containing 80 genes, and n=6, such that at least 6 syntenic genes were required to define a block (the minimum number of anchors in a cluster), and ‘-full’ for predicting reciprocal best hit genes. The ends of a block were defined when its two flanking regions had successive 80 genes which did not show the same syntenic relationship as the block. Thus, each block includes at least 6 syntenic genes separated by 80 genes which do not show the syntenic relationship identified for the block.

To calculate the length of each syntenic block, we used the location information for all genes within the block. and processed these data using BEDTools v2.30.0 with the parameter “mergeBed -d 1800000” to merge adjacent syntenic regions.

### Identification of the loss and reserve genes on chromosome 5

To infer loss and preservation of genes on the *R. natans* chr5, we again used JCVI v1.2.7 with the parameter (--full -n 1) to identify the single best-matching syntenic ‘anchor’ pairs in our three *de novo* genome assemblies. We classified these genes as candidate conserved syntenic genes.

### Sex-related gene identification and phylogenetic analyses

Orthologues of *BPC* and *FGMYB* were obtained from 56 green plant genome data sets, 54 liverworts transcriptomes (Dong et al., 2022), 3 red algae genome sequences (*Chondrus crispus*, Collén et al., 2013; *Porphyra umbilicalis*, Brawley et al., 2017; *Cyanidioschyzon merolae*, Misumi et al., 2005), 1 Prasinodermophyta genome sequence (*Prasinoderma coloniale*, Li et al., 2020), 405 fungal species genome sequences, 24,189 bacterial genome sequences, 496 archaeal species genome sequences, and 11,664 virus genome sequences. The GAGA_bind Pfam model (PF06217) was used in hmmsearch v3.1 (E-value <= 1e-5) (https://www.mankier.com/1/hmmsearch) to search for BPC candidate proteins. Six proteins (CepurGG1.UG291100.1.v1.1, CepurR40.VG015400.1.v1.1, Mp1g17210.1, RF8g03370.1, RNIG4g14090.1, and RN1g16500.1, where Cepur refers to the moss *Ceratodon purpureus* genome sequence protein annotation) were used as query sequences to scan the selected databases with BLASTP (Nucleotide-Nucleotide BLAST 2.12.0+) (E-value <= 30) to identify the candidate *FGMYB* proteins. Four liverworts and the C. *purpureus* genome protein annotations were used as databases to identify the orthologues of other sex-related gene families, including genes specific to the MpchrU and MpchrV, using local BLASTP (E-value <= 1e-30, or E-value <= 1e- 60) searches.

MpchrU- and MpchrV-specific sex chromosome genes described in Iwasaki et al. (Iwasaki et al., 2021) were used, along with protein annotations from four liverwort species and the *C. purpureus* genome, to identify orthologues of the sex chromosome-linked genes, using local BLASTP searches with an E-value threshold of ≤ 1e-30 or ≤ 1e-60.

All candidate sequences were aligned with MUSCLE v5.1.linux64 (Madeira et al., 2022) to generate multiple sequence alignments for each gene family. The alignments were trimmed to delete unconservative regions by trimAl v1.4.rev22 (Capella-Gutiérrez et al., 2009) using automated parameter selection before phylogenetic tree estimation with IQ-TREE 2 v2.2.0.3 (Nguyen et al., 2014) and fasttree v 2.1.11 (Price et al., 2009). All gene tree visualizations were conducted used FigTree v1.4.4 (http://tree.bio.ed.ac.uk/software/figtree/).

## Reporting summary

Further information on research design is available in the Nature Portfolio Reporting Summary linked to this article.

## Data availability

Genome assemblies of three liverwort species and their annotations are available at Figshare (https://doi.org/10.6084/m9.figshare.23805990). All raw sequencing data used for analysis have been deposited at China National GeneBank DataBase (CNGBdb) under project CNP0004629.

## Code availability

Scripts and pipelines applied for analysis in this work can be found freely at GitHub (https://github.com/abysw/Sexual-article2023).

## Acknowledgements

We would like to thank Bengt Hansson at Lund University for critical reading and comments and Brain Charlesworth at University of Edinburgh for insightful discussions. We acknowledge Mengxiao Hu at Hangzhou Botanical Garden, Liguang Sun at Caofeidian Wetland, and Gaojun Liu at Peking University Health Science Center, Shuo Shi at Hebei Normal University, Dechang Meng at South China Botanical Garden CAS, and Zhaojie Ren at Shandong Museum for field collection of liverwort colonies. We thank Herbarium of IBCAS (PE) for providing liverwort specimen. We are grateful to Lijie Zhu at IBCAS for drawing of reproductive structures of liverwort species, Jindan Zhang at IBCAS for assistance with flow cytometer, and Ze Wei at Plant Photo Bank of China for providing liverwort photos. This work is supported by National Natural Science Foundation of China (32221001 to Z.C., Y.J., and B.X. and 32070249 to B.X.), K.C. Wong Education Foundation (GJTD-2020-05, to Z.C., Y.J., and B.X.), and the Special Research Assistant Program of Chinese Academy of Sciences (to X.Z.).

## Author contributions

The project was conceived and designed by F.Y., X.Z., and B.X.. F.Y. and X.Z. performed all experiments unless specified here. T.Z., W.Y., and Y.S. established axenic isolates and cultured plants at various conditions. Q.H. and X.Z. identified liverwort collections. Y.F. X.Z., W.S., J.Z., D.C., Y.J., Z.C., and B.X. analysed data. F.Y., X.Z., D.C., and B.X. wrote the manuscript with input from other authors. All authors read and approved of its content.

## Supplementary Information

**Supplementary Figure S1.** Gametophytes of liverwort species mentioned or investigated in this study.

**Supplementary Figure S2.** Illustration of reproductive structures of the four liverwort species studied.

**Supplementary Figure S3.** Hi-C heat maps of *Ricciocarpos natans* and *Riccia fluitans*.

**Supplementary Figure S4.** Pairwise intra- and inter-species synonymous site divergence estimates (*Ks*) analysis of four liverwort species.

**Supplementary Figure S5.** Gene family analysis of 56 green plant species.

**Supplementary Figure S6.** Analysis of DNA sequence diversity in 50 Kb windows of different chromosomes in two liverwort species.

**Supplementary Figure S7.** Identification of autosomal paralogues of sex-linked genes in *M. polymorpha*.

**Supplementary Figure S8.** Phylogenetic tree of U chromosome-specific gene MpUg00110.

**Supplementary Figure S9.** Phylogenetic trees of U chromosome-specific genes.

**Supplementary Figure S10.** Synteny analysis of the sex chromosome and chromosome 5 in four liverwort species.

**Supplementary Figure S11.** Phylogenetic trees of V chromosome-specific genes.

**Supplementary Figure S12.** Phylogenetic tree of *BPCV/BPCU* genes in green plants.

**Supplementary Figure S13.** Phylogenetic tree of *FGMYB* genes in green plants.

**Supplementary Table S1.** The sexual status of liverworts.

**Supplementary Table S2.** Sequencing data information.

**Supplementary Table S3.** Genome structure.

**Supplementary Table S4.** Gene family analysis.

**Supplementary Table S5.** Fate of sex chromosome-specific genes.

**Supplementary Table S6.** Estimation of chromosome divergence in Marchantiophyta.

