## Supplemental Figures for "Evidence for evolution of a new sex chromosome within the Marchantiales haploid-dominant plant lineage"

*Details*

**Supplementary Figure S1.** Gametophytes of liverwort species mentioned or investigated in this study.

**Supplementary Figure S2.** Illustration of reproductive structures of the four liverwort species studied.

**Supplementary Figure S3.** Hi-C heat maps of *Ricciocarpos natans* and *Riccia fluitans.*

**Supplementary Figure S4.** Histogram of intra- and inter-species synonymous site divergence estimates (*Ks*) values for the four species studied.

**Supplementary Figure S5.** Gene family analysis of 56 green plant species.

**Supplementary Figure S6.** Analysis of DNA sequence diversity in 50 Kb windows of different chromosomes in two liverwort species.

**Supplementary Figure S7.** Identification of autosomal paralogues of sex-linked genes in *M. polymorpha*.

**Supplementary Figure S8.** Phylogenetic tree of U chromosome-specific gene MpUg00110.

**Supplementary Figure S9.** Phylogenetic trees of U chromosome-specific genes.

**Supplementary Figure S10.** Synteny analysis of the sex chromosome and chromosome 5 in four liverwort species.

**Supplementary Figure S11.** Phylogenetic trees of V chromosome-specific genes.

**Supplementary Figure S12.** Phylogenetic tree of *BPCV/BPCU* genes in green plants.

**Supplementary Figure S13.** Phylogenetic tree of *FGMYB* genes in green plants.

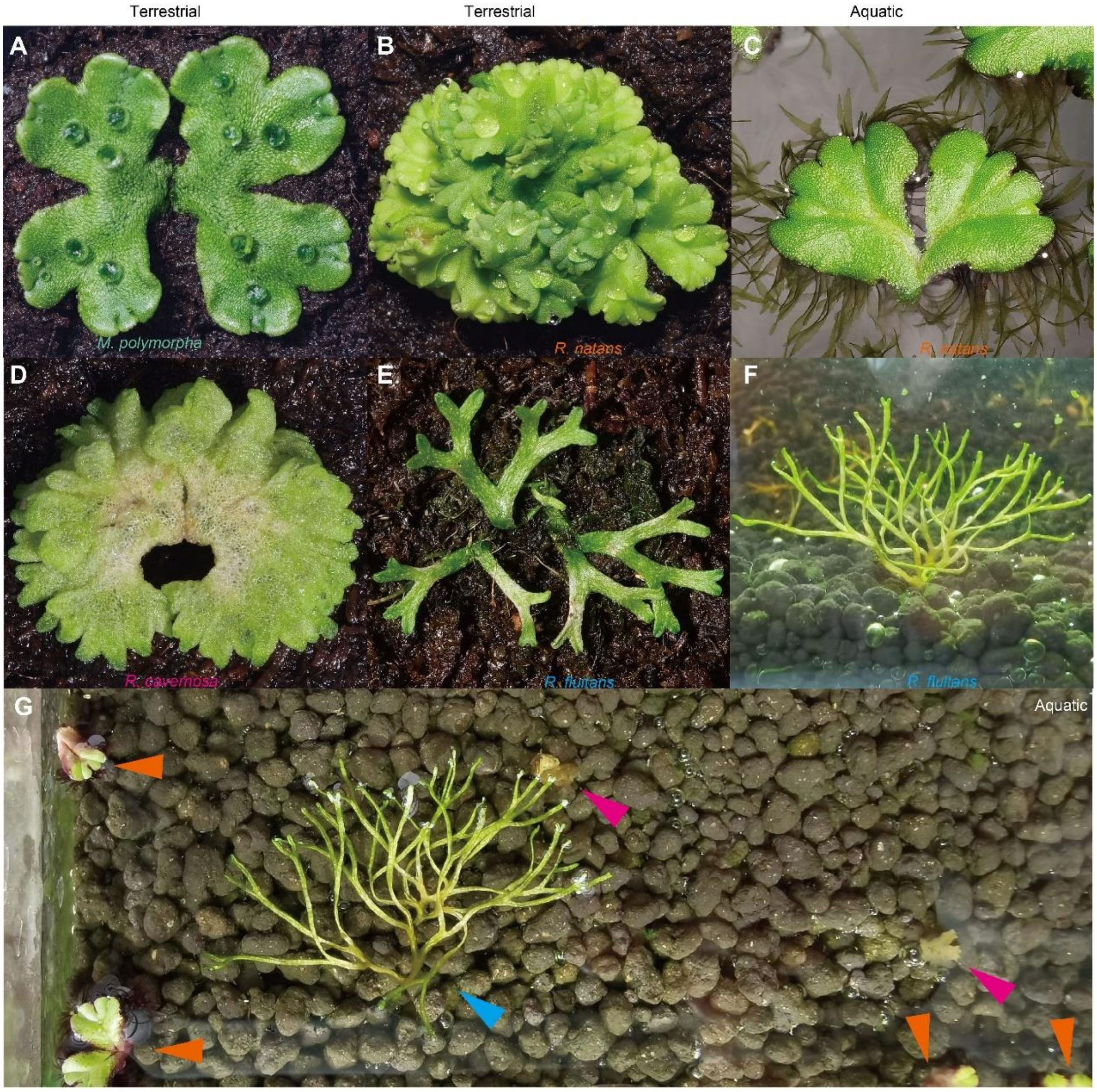


**Supplementary Figure S1.** Gametophytes of liverwort species mentioned or investigated in this study. **A)** *Marchantia polymorpha*. **B) and C)** *Ricciocarpos natans* grown in terrestrial and aquatic habitats. **D)** *Riccia cavernosa*. **E) and F)** *Riccia fluitans* grown in terrestrial and aquatic habitats. **G)** Adaptation of *Ricciocarpos natans* and *Riccia fluitans* to aquatic habitat. Arrow heads in orange, blue, and pink indicate *Ricciocarpos natans*, *Riccia fluitans*, and *Riccia cavernosa* (dead).

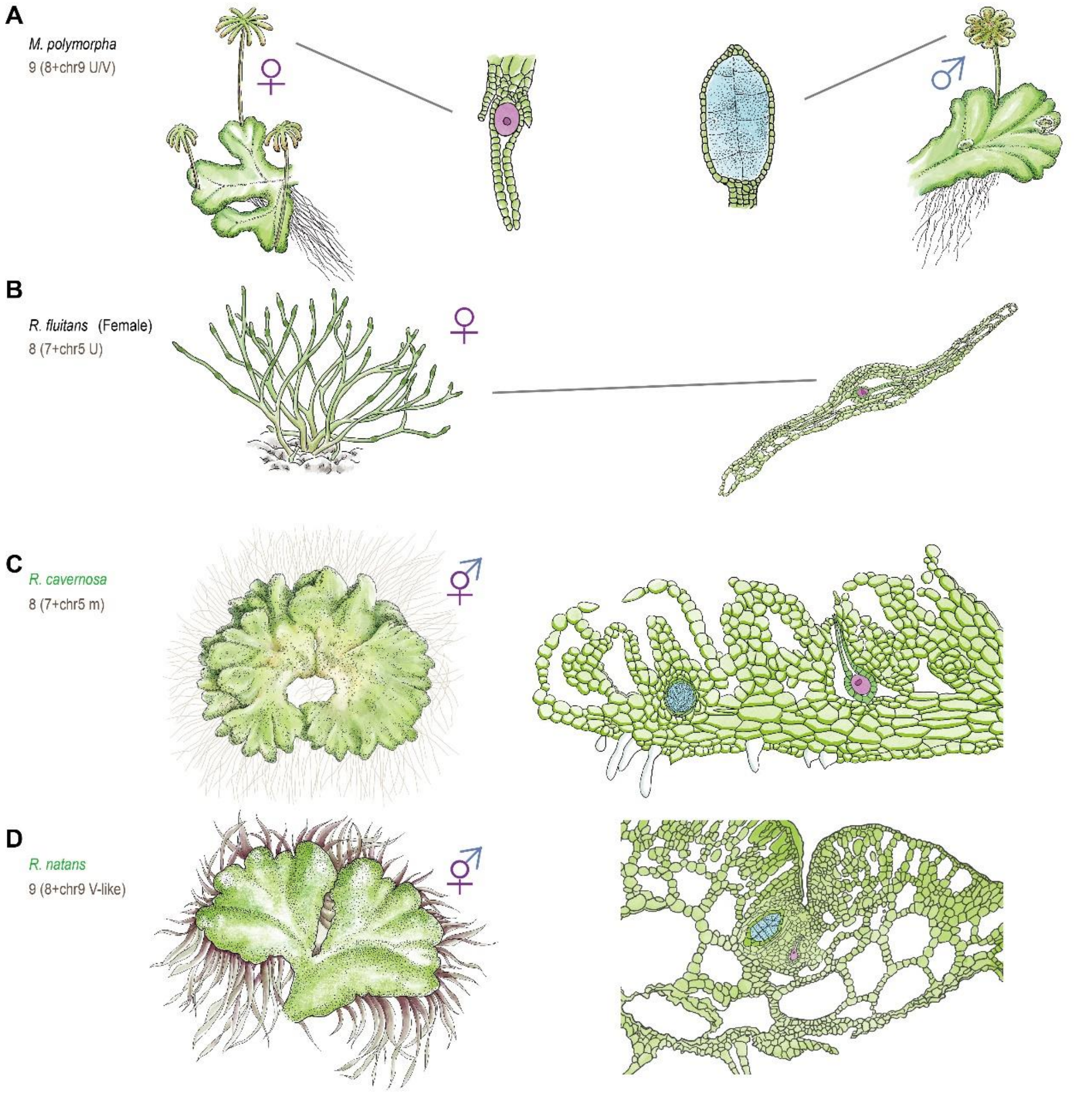


**Supplementary Figure S2.** Illustration of reproductive structures of the four liverwort species studied. **A)** Reproductive organs of a *Riccia fluitans* female, showing archegonia inside the thallus. **B)** Reproductive organs of *Riccia cavernosa*. In the reproductive stage, archegonia and antheridia are inside the thallus. **C)** Reproductive organs of *Riccia natans*. Unlike *M. polymorpha*, no modified reproductive branches develop. Antheridia and archegonia are embedded in the thallus near the midrib. **D)** Reproductive growth of *Marchantia polymorpha*. Female reproductive branches form umbrella-like structures, called archegoniophores (left), bearing archegonia containing egg cells. Male reproductive branches form disc-like structures, called antheridiophores (right), in which antheridia are embedded and produce sperm cells. Female and male germ cells are indicated in red and blue, respectively.

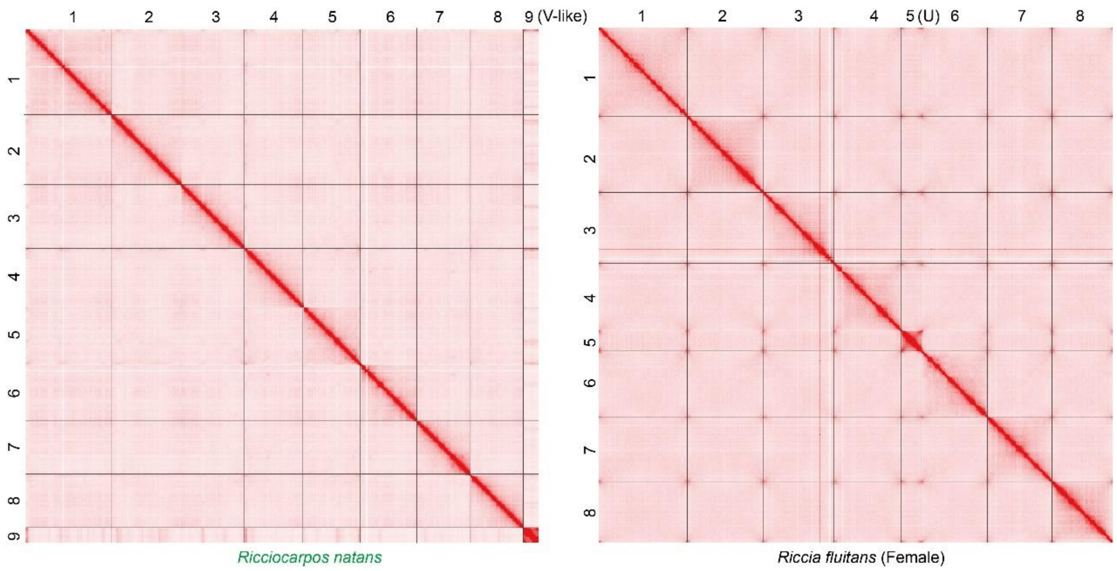


**Supplementary Figure S3.** Hi-C heat maps of *Ricciocarpos natans* and *Riccia fluitans*, showing the unusual results for the *R. fluitans* chromosome 5.

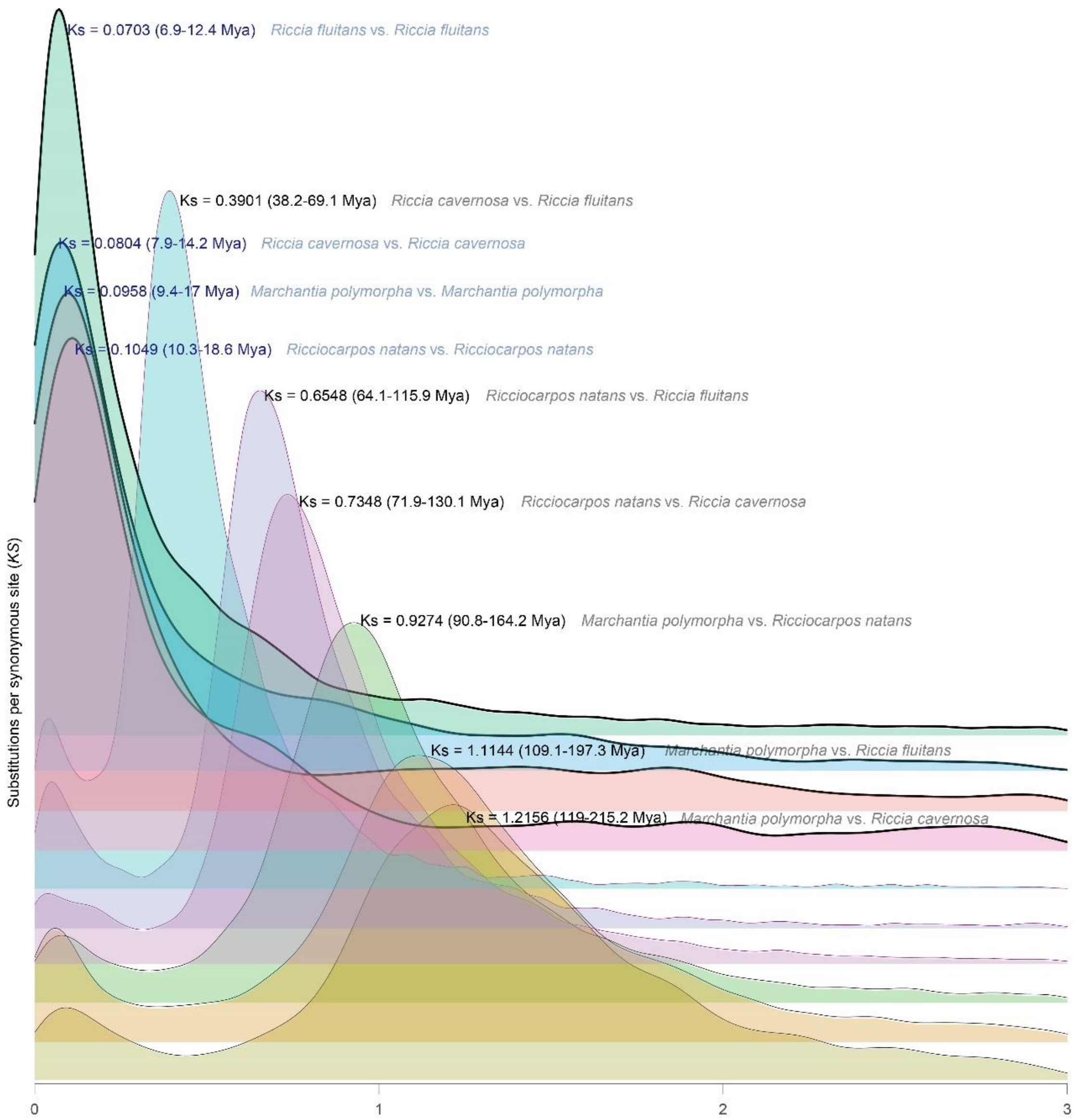


**Supplementary Figure S4.** Histogram of intra- and inter-species synonymous site divergence estimates (*Ks*) values for the four species studied. The three species whose changes in sexual system were studied here, and the outgroup, *M. polymorpha*. The darker line and blue words indicate intra-species, while the lighter line and black/grey words indicate inter-species. The divergence time of 119.0-215.2 Mya between *Marchantia* and *Ricciocarpos* was obtained from the timetree website (https://timetree.org), and is consistent with the time in the study of Villarreal *et al*. (Villarreal et al. 2016)

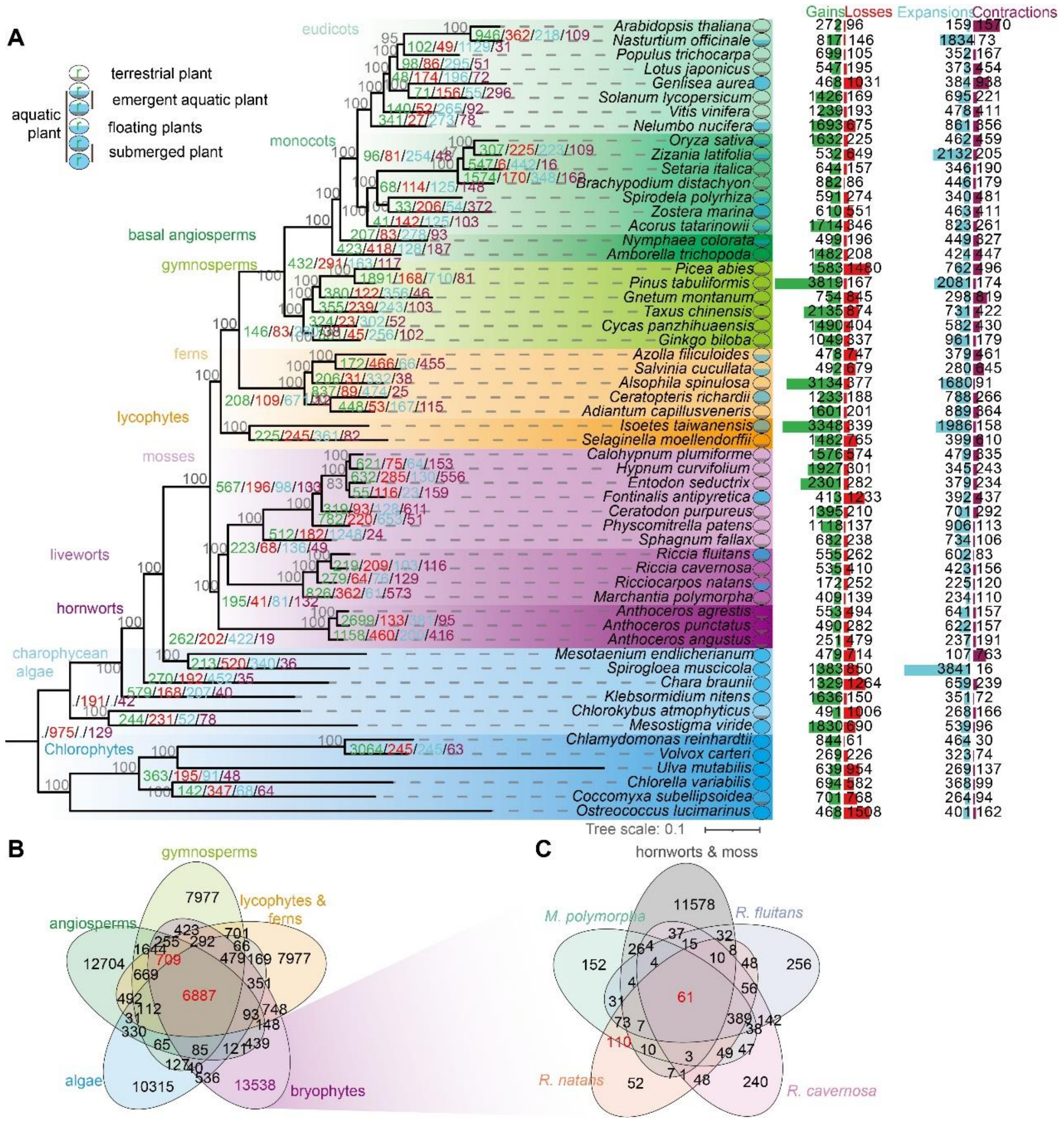


**Supplementary Figure S5.** Gene family analysis of 56 green plant species. **A)** Phylogeny of the 56 species analysed in this study. The evolutionary history of green plants was reconstructed using the inferred amino acid sequences of low copy genes identified from assembled genomes of 56 plant species (see Methods). The colored numbers at each node and the bars with numbers show gains (green), losses (red), expansions (cyan) and contractions (purple) of gene families, respectively. Bootstrap values calculated with 1,000 replicates are indicated in gray at the base of each branch. The scale bar represents the amino acid divergence value per site. **B)** Venn diagram displaying the numbers of gene families shared by all 56 species, based on the analysis explained in the main text. A total of 6,887 gene families are shared in green plants. **C)** Venn diagram of the numbers of gene families shared between the bryophyte species indicated. A total of 61 gene families is shared between the bryophyte species indicated.

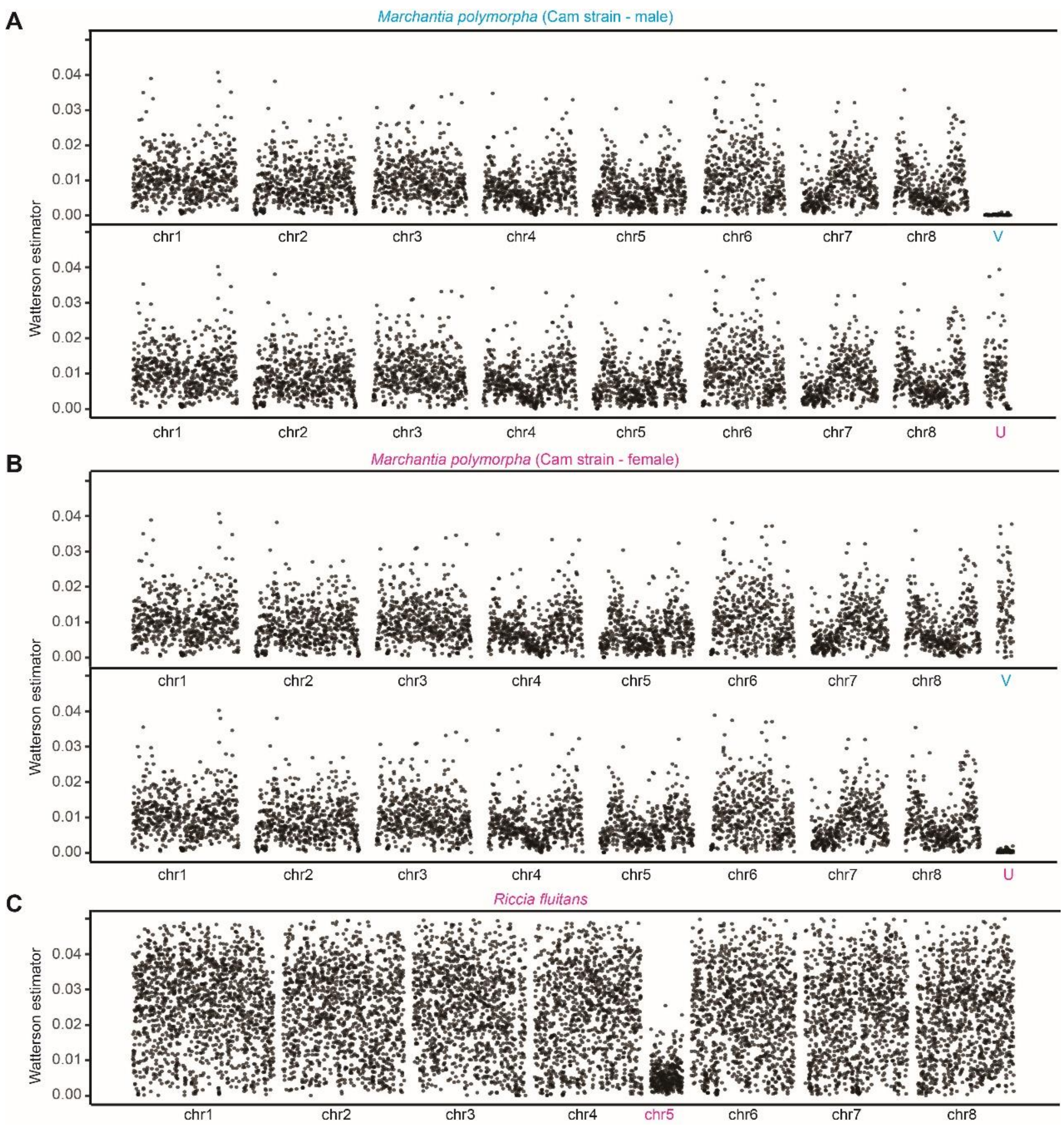


**Supplementary Figure S6.** Analysis of DNA sequence diversity in 50 Kb windows of different chromosomes in two liverwort species. **A) and B)** Pairwise nucleotide diversity estimated from two *Marchantia polymorpha* strains (Tak, the reference genome assembly for this species, and Cam, a sample collected in Cambridge, UK), using males and females. **C)** Nucleotide diversity between four *R. fluitans* colonies collected from geographically distant locations, including Hangzhou (Zhejiang province, China), Shaoguan (Guangdong province, China), Ziyuan (Guangxi province, China), and Alachua (Florida, USA; specimen PE01407550). *R. fluitans* genome (Hangzhou accession) assembled in the present study was used as the reference.

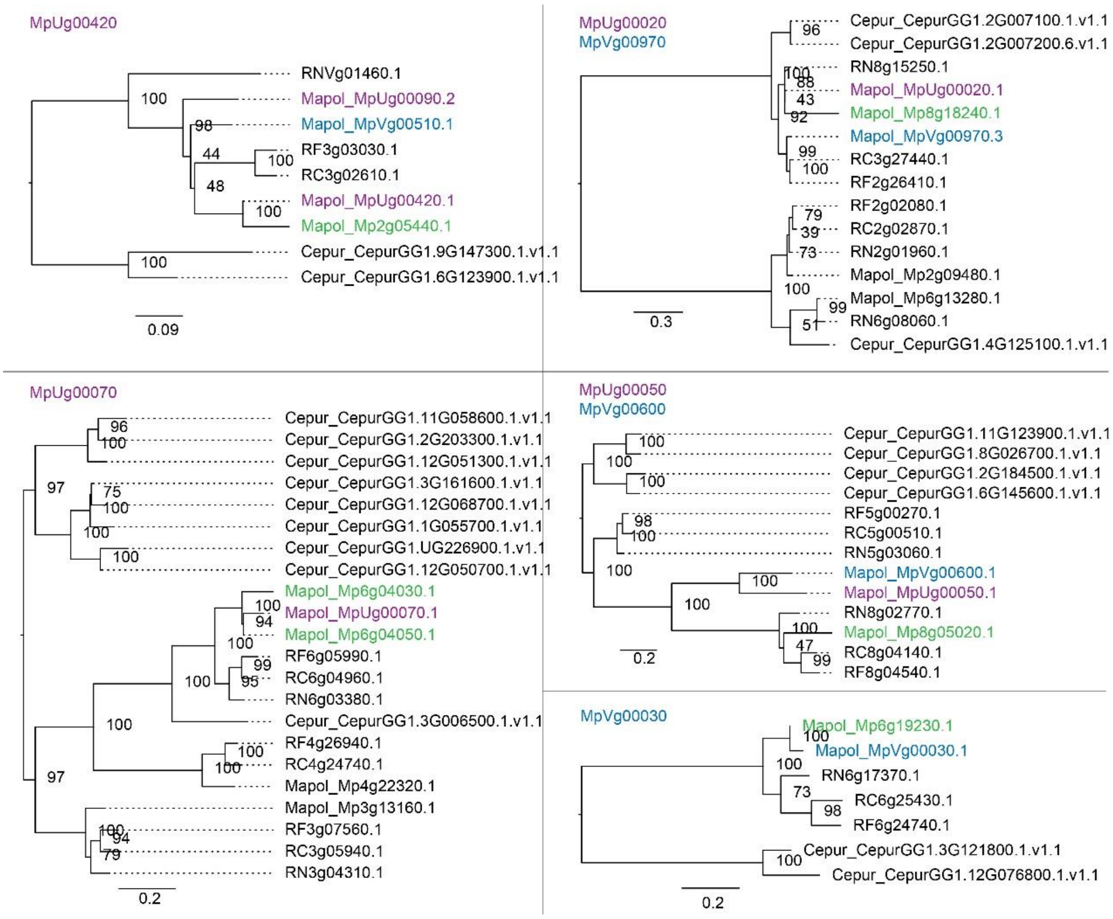


**Supplementary Figure S7.** Identification of autosomal paralogues of sex-linked genes in *M. polymorpha*. Phylogenetic analyses amino acid sequences of U chromosome-specific genes were performed separately for the MpUg00420, MpUg00070, MpUg00020/MpVg00970, MpUg00050/ MpVg00600, MpVg00030 genes, which were defined in ref (Iwasaki et al. 2021). 23 as U- or 39 V-specific genes in *M. polymorpha.* Green font indicates that the *M. polymorpha* genome has an autosomal paralog of the gene indicated in the label for each tree. Numbers at each branch indicate bootstrap values calculated with 1,000 replicates; only values > 50% are only shown. The scale bar represents the amino acid divergence per site. Cepur, *Ceratodon purpureus*; Mapol, *Marchantia polymorpha*; RN, *Ricciocarpos natans*; RC, *Riccia cavernosa;* RF, *Riccia fluitans.*

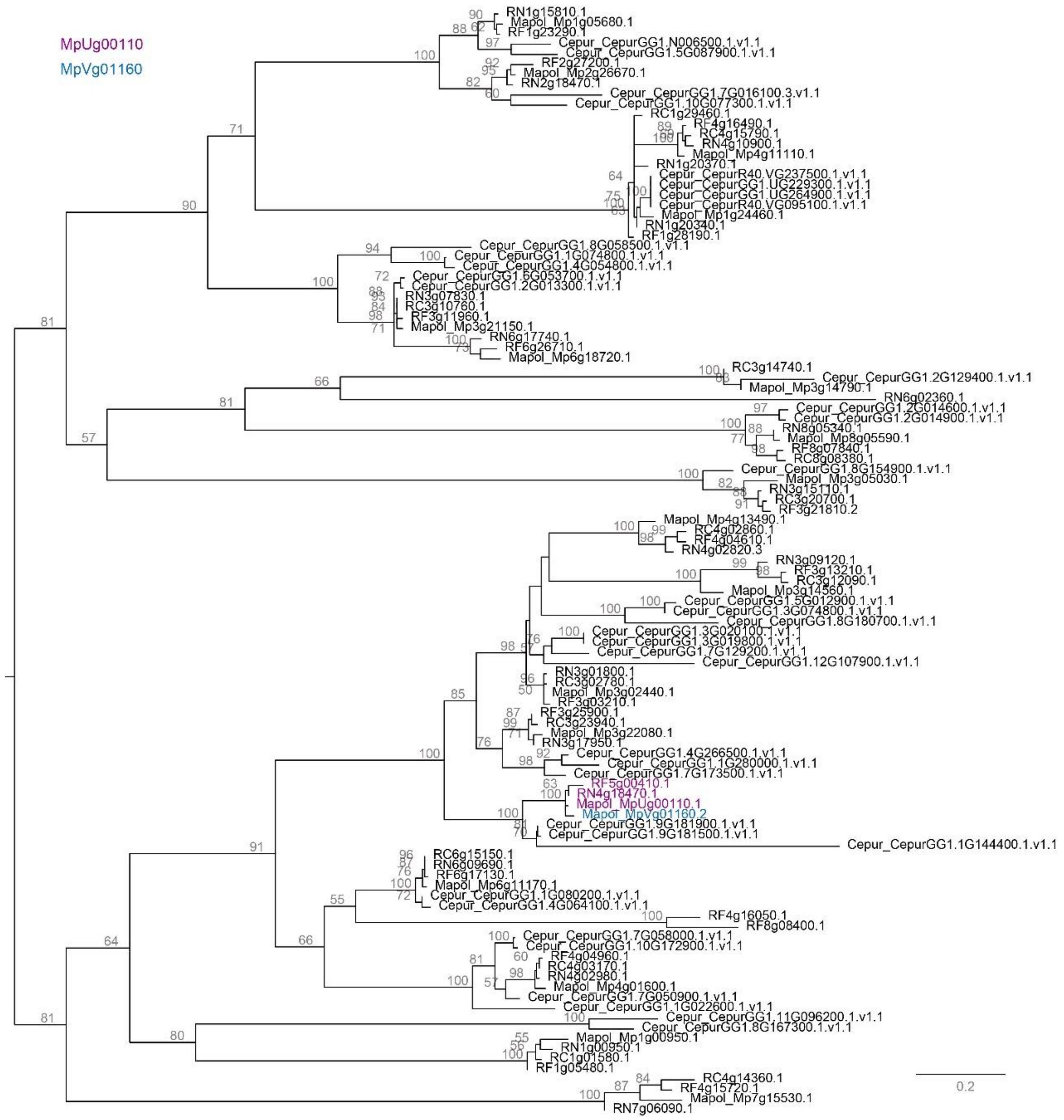


**Supplementary Figure S8.** Phylogenetic tree of U chromosome-specific gene MpUg00110. The phylogeny was constructed with coding sequences homologous to MpUg00110 from five bryophyte species. Numbers at each branch indicate bootstrap values calculated with 1,000 replicates. The bootstrap values more than 50% are only shown. The scale bar represents the amino acid divergence per site. Cepur*, Ceratodon purpureus*; Mapol, *Marchantia polymorpha*; Rinat, *Ricciocarpos natans*; Ricav, *Riccia cavernosa;* Riflu, *Riccia fluitans.*

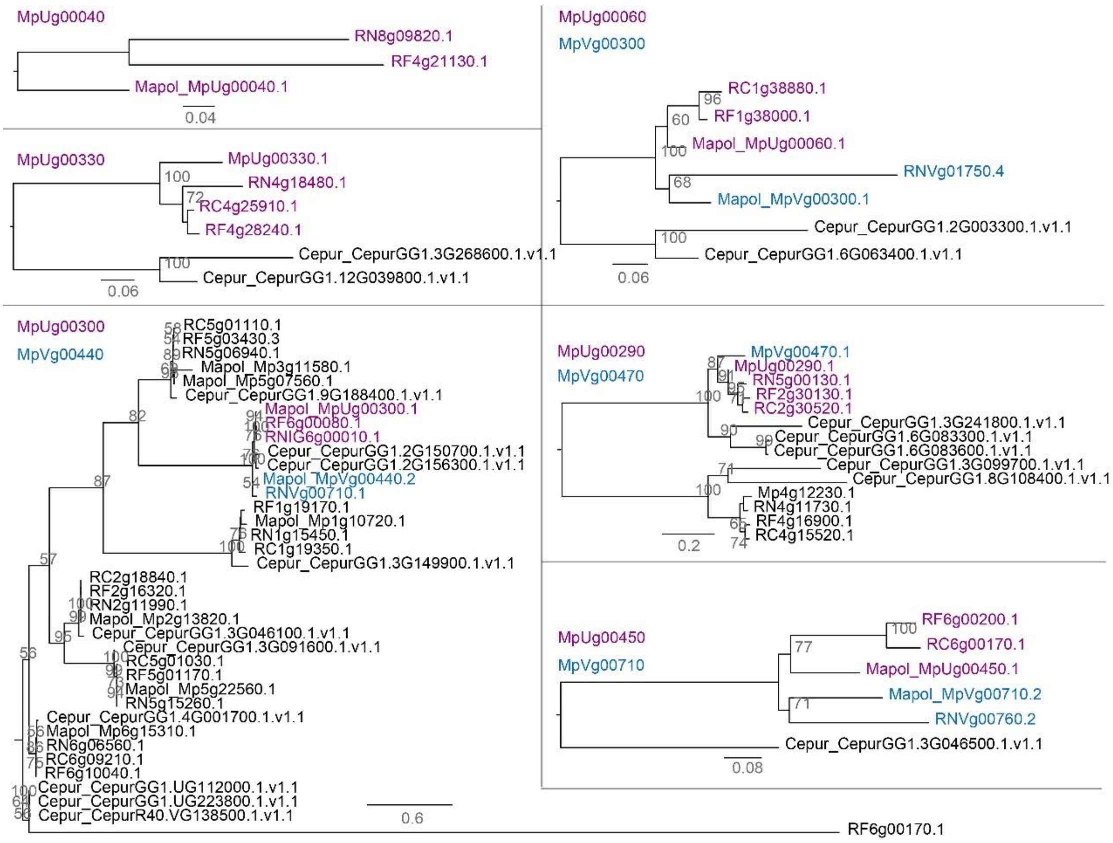


**Supplementary Figure S9.** Phylogenetic trees of U chromosome-specific genes. Phylogenetic analysis of U chromosome-specific genes was performed separately for the MpUg00040, MpUg00060, MpUg00290, MpUg00300, MpUg00330 and MpUg00450 genes*.* Numbers at each branch indicate bootstrap values calculated with 1,000 replicates. The bootstrap values more than 50% are only shown. The scale bar represents the amino acid divergence per site. Cepur*, Ceratodon purpureus*; Mapol, *Marchantia polymorpha*; Rinat, *Ricciocarpos natans*; Ricav, *Riccia cavernosa;* Riflu, *Riccia fluitans.*

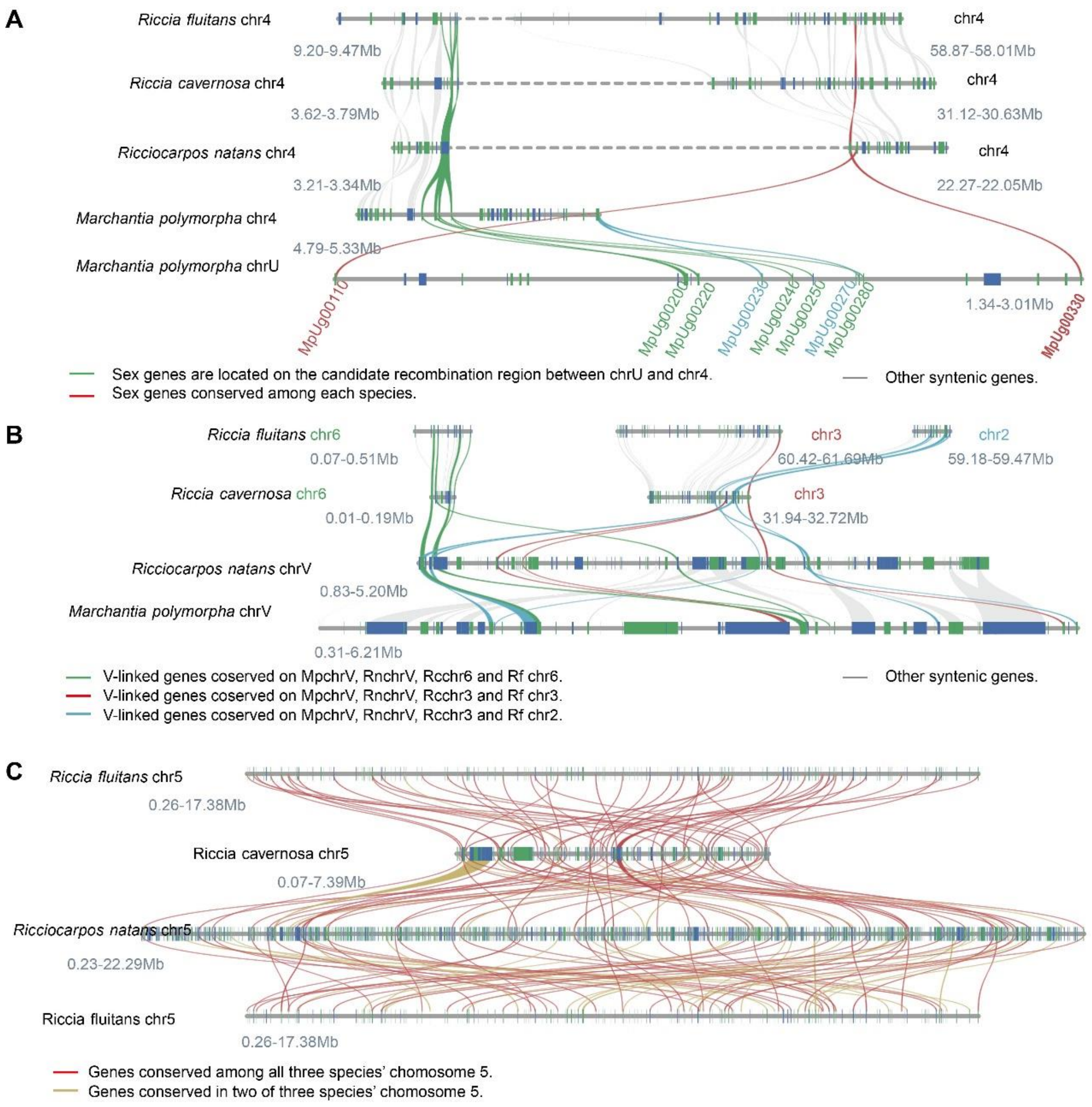


**Supplementary Figure S10.** Synteny analysis of the sex chromosome and chromosome 5 in four liverwort species. **A)** Analysis of segments of the *M. polymorpha* U chromosome and homologous regions of chr4 in *M. polymorpha* and the three other liverworts. **B)** Analysis of segments of the V chromosome of *Marchantia* and autosomes of *Ricciocarpos* and *Riccia*. **C)** Analysis of chromosome 5 of different Ricciaceae species. Note the low gene density on chromosome 5 compared with the close outgroup, *R. natans*. The top and bottom chromosome diagrams show the *R. fluitans* chromosome 5, and the middle two show the homologous chromosome in the two other species. Vertical bars on chromosome diagrams represent gene loci, and gene orientations are indicated in blue (forward orientation) and green (reverse orientation) respectively.

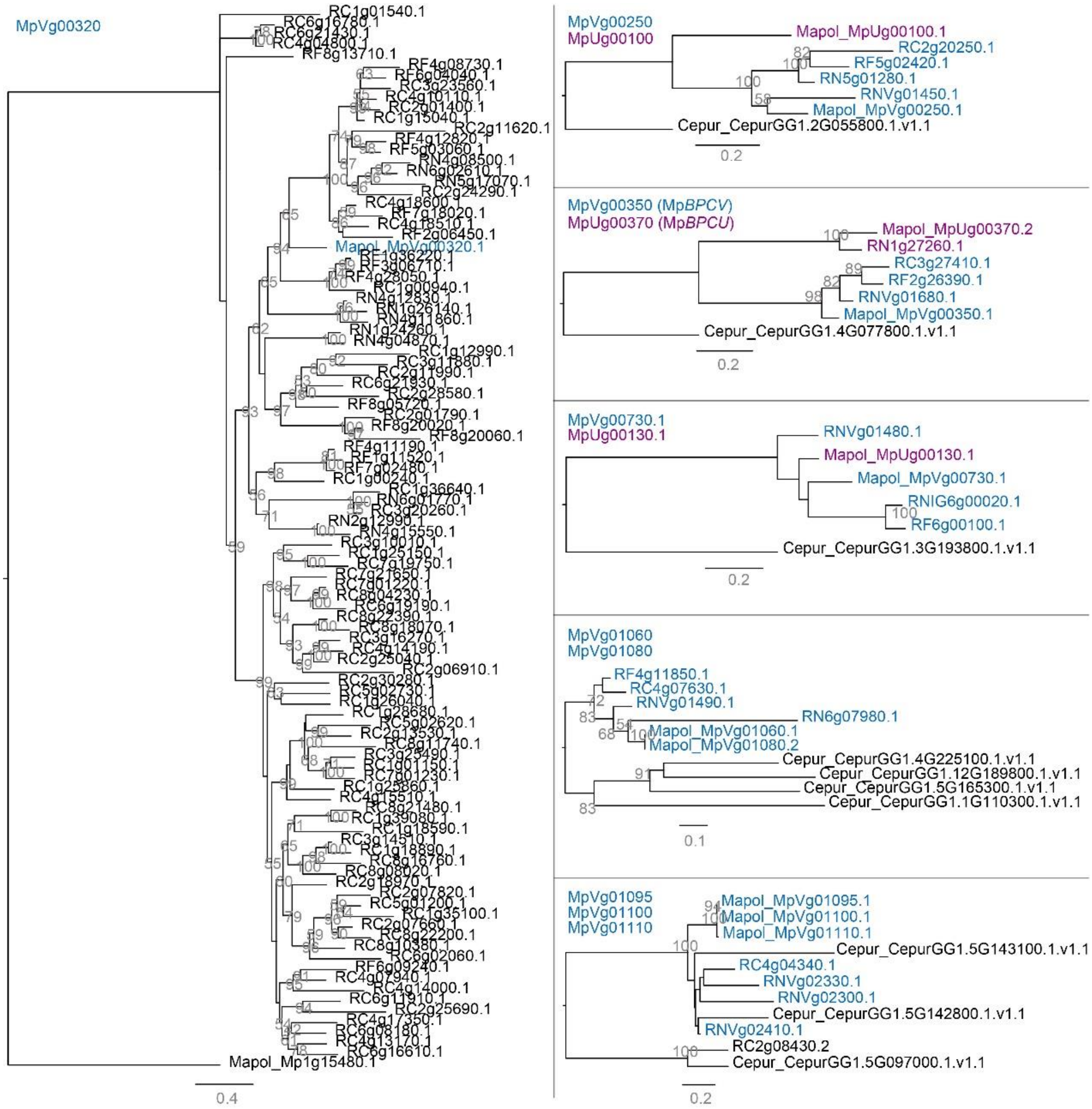

**Supplementary Figure S11.** Phylogenetic trees of V chromosome-specific genes. Phylogenetic analysis of U chromosome-specific genes, performed separately for the MpVg00320, MpVg00250, MpVg00350, MpVg00730, MpVg01060, MpVg01080, MpVg01095, MpVg01100, MpVg01110 genes. Numbers at each branch indicate bootstrap values calculated with 1,000 replicates. The bootstrap values more than 50% are only shown. The scale bar represents the amino acid divergence per site. Cepur*, Ceratodon purpureus*; Mapol, *Marchantia polymorpha*; Rinat, *Ricciocarpos natans*; Ricav, *Riccia cavernosa;* Riflu, *Riccia fluitans.*

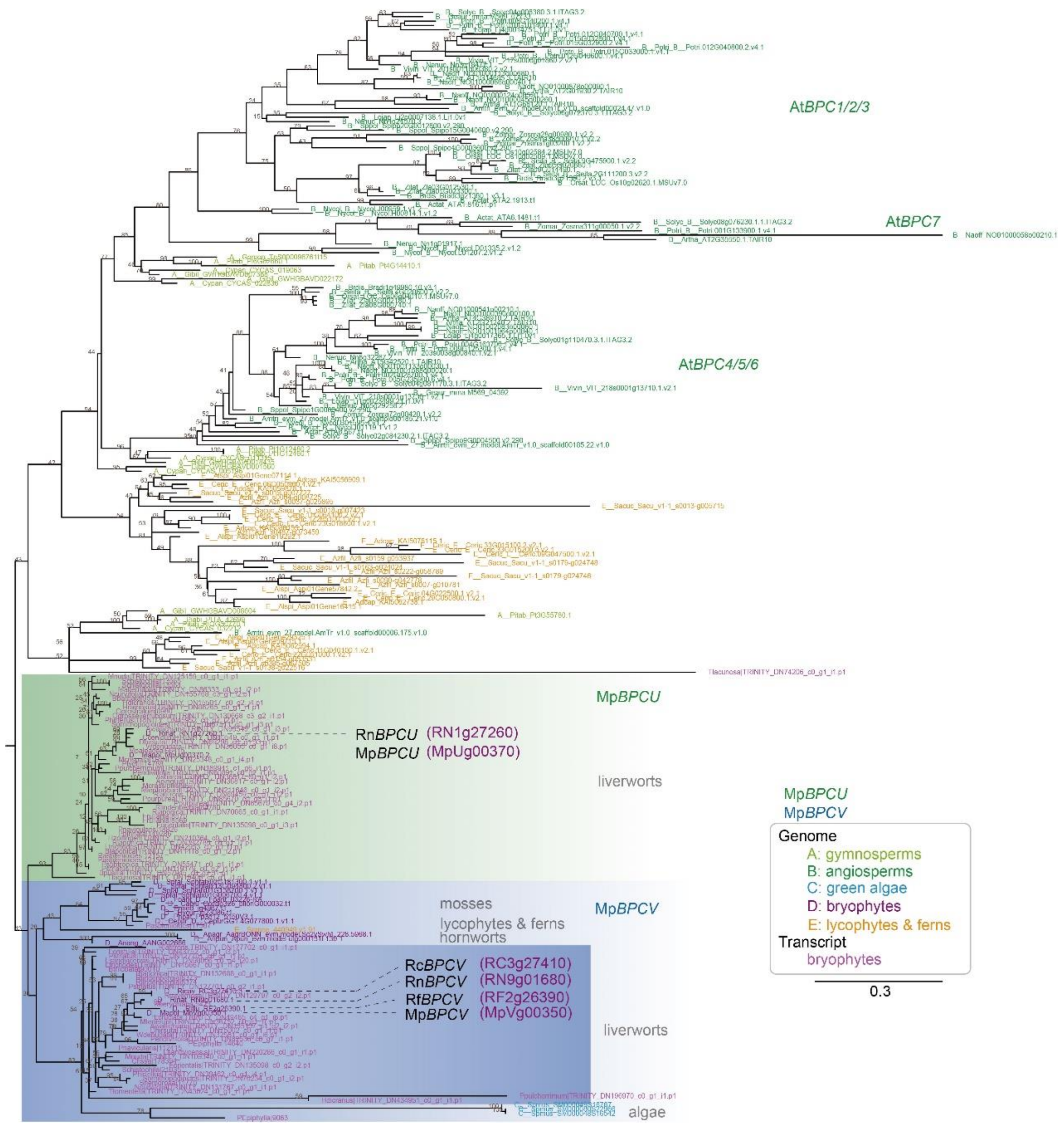


**Supplementary Figure S12.** Phylogenetic tree of *BPCV/BPCU* genes in green plants. A total of 146 coding sequences of *FGMYB* genes were identified from 110 representative plant species (56 genome sequences and 54 transcriptome data sets). Numbers at each branch indicate bootstrap values calculated with 1,000 replicates. The bootstrap values more than 50% are only shown. The scale bar represents the amino acid divergence per site. Anpin, *Aneura pinguis*; Ansha, *Aneura sharpi*i; Aswal, *Asterella wallichiana*; Babar, *Barbilophozia barbata*; Bajap, *Bazzania japonica*; Batri, *Bazzania trilobata*; Blpus, *Blasia pusilla*; Cafis, *Calypogeia fissa*; Cocon, *Conocephalum conicum*; Duhir, *Dumortiera hirsuta*; Focri, *Fossombronia cristula*; Frori, *Frullania orientalis*; Frori, *Frullania orientalis*; Hamni, *Haplomitrium mnioides*; Hedic, *Herbertus dicranus*; Heram, *Herbertus ramosus*; Hezol, *Heteroscyphus zollingeri*; Lesan, *Lepidozia sandvicensis*; Letri, *Lepidozia trichodes*; Lucru, *Lunularia cruciata*; Macri, *Makinoa crispata*; Mapal, *Marchantia paleacea*; Mapol, *Marchantia polymorpha*; Mealt, *Metacalypogeia alternifolia*; Mecra,

*Metzgeria crassipilis*; Melep, *Metzgeria leptoneura*; Moten, *Monosolenium tenerum*; Mynud, *Mylia nuda*; Nocur, *Nowellia curvifolia*; Odgro, *Odontoschisma grosseverrucosum*; Odpro, *Odontoschisma prostratum*; Palye, *Pallavicinia lyellii*; Peend, *Pellia endiviifolia*; Peepi, *Pellia epiphylla*; Plasp, *Plagiochila asplenioides*; Plsub, *Plagiochila subtropica*; Plpur, *Pleurozia purpurea*; Plhir, *Plicanthus hirtellus*; Ponav, *Porella navicularis*; Popin, *Porella pinnata*; Poplu, *Porella plumosa*; Ptpul, *Ptilidium pulcherrimum*; Ptstr, *Ptychanthus striatus*; Rajap, *Radula japonica*; Ralin, *Radula lindenbergia*; Rilat, *Riccardia latifrons*; Ricav, *Riccia cavernosa*; Riflu, *Riccia fluitans*; Rinat, *Ricciocarpos natans*; Scnem, *Scapania nemorosa*; Scorn, *Scapania ornithopodioides*; Scorn, *Scapania ornithopodioides*; Sptex, *Sphaerocarpos texanus*; Trlac, *Treubia lacunosa*; Trtom, *Trichocolea tomentella*; Widen, *Wiesnerella denudata*. For transcriptomes: Apinguis, *Aneura pinguis*; Asharpii, *Aneura sharpii*; Awallichiana, *Asterella wallichiana*; Btrilobata, *Barbilophozia barbata*; Bjaponica, *Bazzania japonica*; Btrilobata, *Bazzania trilobata*; Bpusilla, *Blasia pusilla*; Cfissa, *Calypogeia fissa*; Cconicum, *Conocephalum conicum*; Dhirsuta, *Dumortiera hirsuta*; Fcristula, *Fossombronia cristula*; Forientalis, *Frullania orientalis*; Frullania, *Frullania sp.*; Hmnioides, *Haplomitrium mnioides*; Hdicranus, *Herbertus dicranus*; Hramosus, *Herbertus ramosus*; Hzollingeri, *Heteroscyphus zollingeri*; Lsandvicensis, *Lepidozia sandvicensis*; Ltrichodes, *Lepidozia trichodes*; Lcruciata, *Lunularia cruciata*; Mcrispata, *Makinoa crispata*; Mpaleacea, *Marchantia paleacea*; Malternifolia, *Metacalypogeia alternifolia*; Mcrassipilis, *Metzgeria crassipilis*; Mleptoneura, *Metzgeria leptoneura*; Mtenerum, *Monosolenium tenerum*; Mnuda, *Mylia nuda*; Ncurvifolia, *Nowellia curvifolia*; Ogrosseverrucosum, *Odontoschisma grosseverrucosum*; Oprostratum, *Odontoschisma prostratum*; Plyellii, *Pallavicinia lyellii*; PEpiphylla, *Pellia cf epiphylla*; Pendiviifolia, *Pellia endiviifolia*; Pasplenioides, *Plagiochila asplenioides*; Psubtropica, *Plagiochila subtropica*; Ppurpurea, *Pleurozia purpurea*; Phirtellus, *Plicanthus hirtellus*; Pnavicularis, *Porella navicularis*; Ppinnata, *Porella pinnata*; Pplumosa, *Porella plumosa*; Ppulcherrimum, *Ptilidium pulcherrimum*; Pstriatus, *Ptychanthus striatus*; Rjaponica, *Radula japonica*; Rlindenbergia, *Radula lindenbergia*; Rlatifrons, *Riccardia latifrons*; Rberychiana, *Riccia berychiana*; Rcavernosa, *Riccia cavernosa*; Snemorosa, *Scapania nemorosa*; Sornithopodioides, *Scapania ornithopodioides*; Schistochila, *Schistochila sp.*; Stexanus, *Sphaerocarpos texanus*; Tlacunosa, *Treubia lacunosa*; Ttomentella, *Trichocolea tomentella*; Wdenudata, *Wiesnerella denudate*.

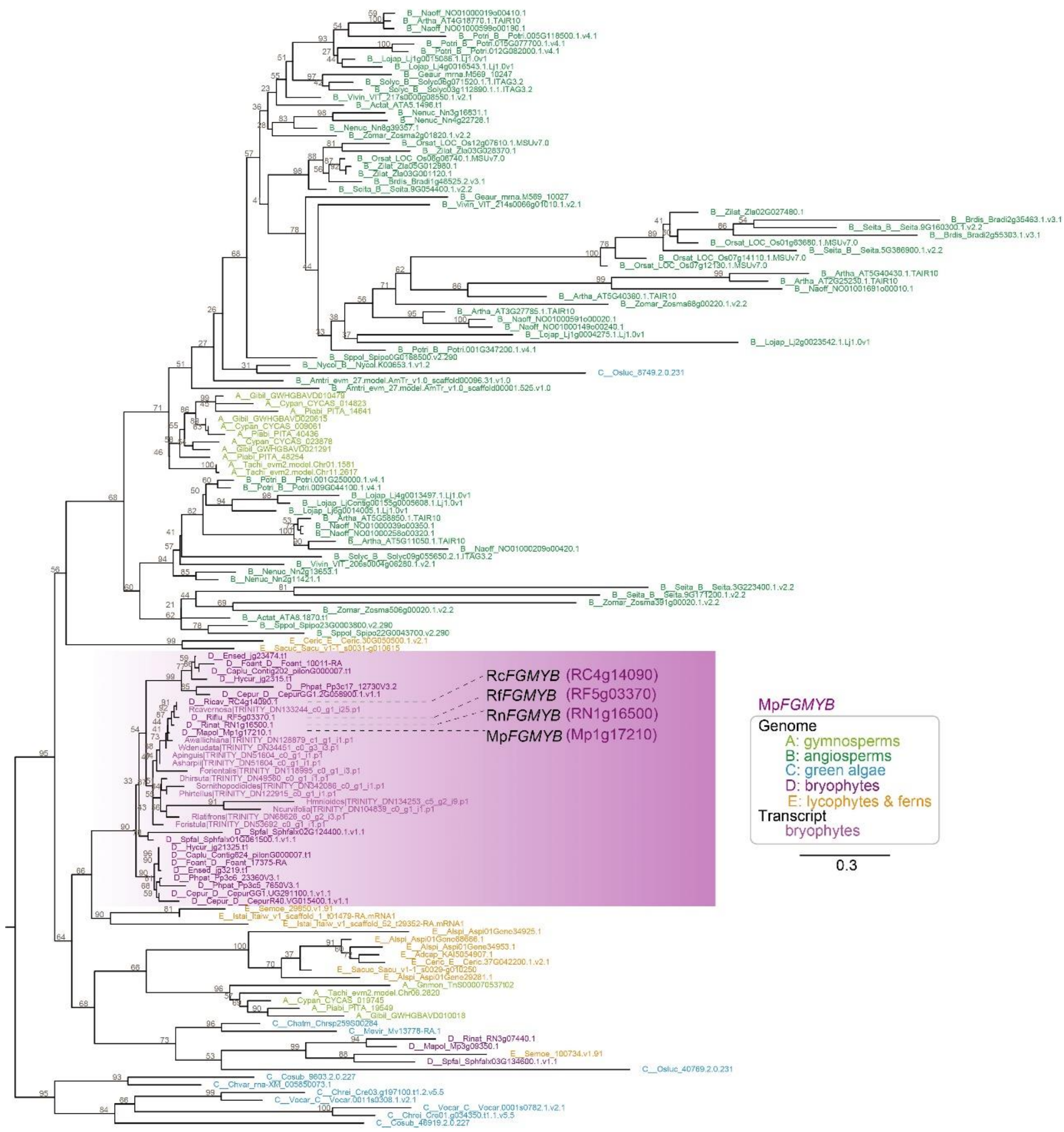


**Supplementary Figure S13.** Phylogenetic tree of *FGMYB* genes in green plants. A total of 229 coding sequences of *FGMYB* genes were identified from 120 representative plant species (56 genome sequences and 54 transcriptome data sets). Numbers at each branch indicate bootstrap values calculated with 1,000 replicates. The bootstrap values more than 50% are only shown. The scale bar represents the amino acid divergence per site. Anpin, *Aneura pinguis*; Ansha, *Aneura sharpii*; Aswal, *Asterella wallichiana*; Babar, *Barbilophozia barbata*; Bajap, *Bazzania japonica*; Batri, *Bazzania trilobata*; Blpus, *Blasia pusilla*; Cafis, *Calypogeia fissa*; Cocon, *Conocephalum conicum*; Duhir, *Dumortiera hirsuta*; Focri, *Fossombronia cristula*; Frori, *Frullania orientalis*; Frori, *Frullania orientalis*; Hamni, *Haplomitrium mnioides*; Hedic, *Herbertus dicranus*; Heram, *Herbertus ramosus*; Hezol, *Heteroscyphus zollingeri*; Lesan, *Lepidozia sandvicensis*; Letri, *Lepidozia trichodes*; Lucru, *Lunularia cruciata*; Macri, *Makinoa crispata*; Mapal, *Marchantia paleacea*; Mapol, *Marchantia polymorpha*; Mealt, *Metacalypogeia alternifolia*; Mecra,

*Metzgeria crassipilis*; Melep, *Metzgeria leptoneura*; Moten, *Monosolenium tenerum*; Mynud, *Mylia nuda*; Nocur, *Nowellia curvifolia*; Odgro, *Odontoschisma grosseverrucosum*; Odpro, *Odontoschisma prostratum*; Palye, *Pallavicinia lyellii*; Peend, *Pellia endiviifolia*; Peepi, *Pellia epiphylla*; Plasp, *Plagiochila asplenioides*; Plsub, *Plagiochila subtropica*; Plpur, *Pleurozia purpurea*; Plhir, *Plicanthus hirtellus*; Ponav, *Porella navicularis*; Popin, *Porella pinnata*; Poplu, *Porella plumosa*; Ptpul, *Ptilidium pulcherrimum*; Ptstr, *Ptychanthus striatus*; Rajap, *Radula japonica*; Ralin, *Radula lindenbergia*; Rilat, *Riccardia latifrons*; Ricav, *Riccia cavernosa*; Riflu, *Riccia fluitans*; Rinat, *Ricciocarpos natans*; Scnem, *Scapania nemorosa*; Scorn, *Scapania ornithopodioides*; Scorn, *Scapania ornithopodioides*; Sptex, *Sphaerocarpos texanus*; Trlac, *Treubia lacunosa*; Trtom, *Trichocolea tomentella*; Widen, *Wiesnerella denudata*. For transcriptomes: Apinguis, *Aneura pinguis*; Asharpii, *Aneura sharpii*; Awallichiana, *Asterella wallichiana*; Btrilobata, *Barbilophozia barbata*; Bjaponica, *Bazzania japonica*; Btrilobata, *Bazzania trilobata*; Bpusilla, *Blasia pusilla*; Cfissa, *Calypogeia fissa*; Cconicum, *Conocephalum conicum*; Dhirsuta, *Dumortiera hirsuta*; Fcristula, *Fossombronia cristula*; Forientalis, *Frullania orientalis*; Frullania, *Frullania sp.*; Hmnioides, *Haplomitrium mnioides*; Hdicranus, *Herbertus dicranus*; Hramosus, *Herbertus ramosus*; Hzollingeri, *Heteroscyphus zollingeri*; Lsandvicensis, *Lepidozia sandvicensis*; Ltrichodes, *Lepidozia trichodes*; Lcruciata, *Lunularia cruciata*; Mcrispata, *Makinoa crispata*; Mpaleacea, *Marchantia paleacea*; Malternifolia, *Metacalypogeia alternifolia*; Mcrassipilis, *Metzgeria crassipilis*; Mleptoneura, *Metzgeria leptoneura*; Mtenerum, *Monosolenium tenerum*; Mnuda, *Mylia nuda*; Ncurvifolia, *Nowellia curvifolia*; Ogrosseverrucosum, *Odontoschisma grosseverrucosum*; Oprostratum, *Odontoschisma prostratum*; Plyellii, *Pallavicinia lyellii*; PEpiphylla, *Pellia cf epiphylla*; Pendiviifolia, *Pellia endiviifolia*; Pasplenioides, *Plagiochila asplenioides*; Psubtropica, *Plagiochila subtropica*; Ppurpurea, *Pleurozia purpurea*; Phirtellus, *Plicanthus hirtellus*; Pnavicularis, *Porella navicularis*; Ppinnata, *Porella pinnata*; Pplumosa, *Porella plumosa*; Ppulcherrimum, *Ptilidium pulcherrimum*; Pstriatus, *Ptychanthus striatus*; Rjaponica, *Radula japonica*; Rlindenbergia, *Radula lindenbergia*; Rlatifrons, *Riccardia latifrons*; Rberychiana, *Riccia berychiana*; Rcavernosa, *Riccia cavernosa*; Snemorosa, *Scapania nemorosa*; Sornithopodioides, *Scapania ornithopodioides*; Schistochila, *Schistochila sp.*; Stexanus, *Sphaerocarpos texanus*; Tlacunosa, *Treubia lacunosa*; Ttomentella, *Trichocolea tomentella*; Wdenudata, *Wiesnerella denudate*.
